# Cross-platform transcriptional profiling identifies common and distinct molecular pathologies in Lewy Body diseases

**DOI:** 10.1101/2021.04.22.440800

**Authors:** Rahel Feleke, Regina H. Reynolds, Amy Smith, Bension Tilley, Sarah A. Gagliano Taliun, John Hardy, Paul M. Matthews, Steve Gentleman, David Owen, Michael R. Johnson, Prashant Srivastava, Mina Ryten

**Affiliations:** Department of Brain Sciences, Imperial College London, London, UK; Department of Neurodegenerative Disease, University College London; Great Ormond Street Institute of Child Health, Genetics and Genomic Medicine, University College London, London, UK; NIHR Great Ormond Street Hospital Biomedical Research Centre, University College London, London, UK; UK Dementia Research Institute at Imperial College London, London, UK; Department of Medicine & Department of Neurosciences, Université de Montréal, Montréal, Québec, Canada; Montréal Heart Institute, Montréal, Québec, Canada; UK Dementia Research Institute at University College London, London, UK; National Heart & Lung Institute, Imperial College London, London, UK

## Abstract

Parkinson’s disease (PD), Parkinson’s disease with dementia (PDD) and dementia with Lewy bodies (DLB) are three clinically, genetically and neuropathologically overlapping neurodegenerative diseases collectively known as the Lewy body diseases (LBDs). A variety of molecular mechanisms have been implicated in PD pathogenesis, but the mechanisms underlying PDD and DLB remain largely unknown, a knowledge gap that presents an impediment to the discovery of disease-modifying therapies. Transcriptomic profiling can contribute to addressing this gap, but remains limited in the LBDs. Here, we applied paired bulk-tissue and single-nucleus RNA-sequencing to anterior cingulate cortex samples derived from 28 individuals, including healthy controls, PD, PDD and DLB cases (n = 7 per group), to transcriptomically profile the LBDs. Using this approach, we (i) found transcriptional alterations in multiple cell types across the LBDs; (ii) discovered evidence for widespread dysregulation of RNA splicing, particularly in PDD and DLB; (iii) identified potential splicing factors, with links to other dementia-related neurodegenerative diseases, coordinating this dysregulation; and (iv) identified transcriptomic commonalities and distinctions between the LBDs that inform understanding of the relationships between these three clinical disorders. Together, these findings have important implications for the design of RNA-targeted therapies for these diseases and highlight a potential molecular “window” of therapeutic opportunity between the initial onset of PD and subsequent development of Lewy body dementia.

## Introduction

The Lewy body diseases (LBDs) comprise three neurodegenerative diseases, which are characterised by accumulation of Lewy bodies (α-synuclein-containing aggregates) in neurons and neuronal processes [1,2]. These disorders, which include Parkinson’s disease (PD), Parkinson’s disease with dementia (PDD) and dementia with Lewy bodies (DLB), have a prevalence in the general population aged ≥ 65 years of 2-3% [3], 0.3-0.5% [4] and 1-2% [1], respectively. Together, PDD and DLB are collectively known as the Lewy body dementias and they are second only to Alzheimer’s disease (AD) in prevalence among people with dementia[5]. All three LBDs are associated with disability and reduced quality of life; DLB is associated with earlier mortality and a higher cost of care compared with AD [6–8]. With no disease-modifying therapies available for any of the LBDs, these diseases present a major unmet clinical need [9].

While a variety of mechanisms, including mitochondrial and lysosomal dysfunction, oxidative stress, α-synuclein misfolding and neuroinflammation, have been implicated in PD pathogenesis [3,10], less is known about the mechanisms underlying PDD and DLB. Elucidating these mechanisms could provide a biological basis for the clinical distinction between PDD and DLB, which remains controversial in the field [1,11–14]. Clinically, PDD and DLB are arbitrarily separated by the diagnostic “1-year rule”: if dementia is diagnosed before or within one year of the onset of parkinsonism, it is considered to represent DLB, whereas PDD is defined by dementia first presenting more than one year after the onset of parkinsonism [15,16]. Thus, PDD and DLB are clinically distinguished based only on the relative timing of motor and cognitive impairments, despite sharing many symptoms (e.g. dementia, depression, parkinsonism, REM sleep behaviour disorder and visual hallucinations). Arguably, two of the core clinical features of DLB, fluctuating cognition and visual hallucinations, are more prevalent in DLB compared with PD/PDD [17,18], suggesting two separate disorders. However, the overlap of these core clinical features could also be evidence that the disorders are on a spectrum of disease, where DLB represents a more severe form of PDD.

Neuropathologically, all three LBDs are classed as synucleinopathies, but at the end stage of disease they often present with concomitant pathologies such as tau neurofibrillary tangles and amyloid-β [19–21]. It has been argued that PDD and DLB can be neuropathologically distinguished from PD on the basis of (i) Lewy body pathology extending beyond the brainstem to limbic and neocortical regions, (ii) a higher α-synuclein load, and (iii) tau and amyloid-β pathology at a more advanced stage [15,16,21]. However, while neuropathological differences have been reported, the extent to which they permit confident distinction between the LBDs when no clinical diagnosis is present remains contentious. Genetically, the differences between PDD and DLB are not well-characterised, although *APOE*, *GBA* and *SNCA* mutations have been implicated in both [14,22]. More is known about the genetic risk factors contributing to PD and DLB, which share some risk loci (*GBA*, *TMEM175* and *SNCA*) and pathways (lysosomal and endocytic pathways) [23–27]. However, there is also evidence that association signals at *SNCA* may be distinct in PD and DLB (i.e. located at the 3’ and 5’ end of the *SNCA* gene, respectively) [23,24,26,28], and while risk pathways are shared, PD genetic risk factors only explain a small portion of DLB phenotypic variance [26,29].

Identifying therapeutic targets that could modify the development of PDD or DLB requires an understanding of the cellular and molecular features of these diseases. Transcriptomic profiling, through RNA-sequencing of patient-derived tissue, would aid in the identification of such targets, but remains limited in all three LBDs. Of all transcriptomic studies of PD and Lewy body dementia highlighted in two recent systematic reviews (33 and 31 gene expression studies in brain, respectively [10,30]), only 5 used RNA-sequencing. Furthermore, among transcriptomic studies of the three LBDs, few have addressed possible alternative splicing or the confounding of bulk-tissue transcriptomic profiling by differences in cellular composition.

Here, we pair bulk-tissue and single-nucleus RNA-sequencing to gain a comprehensive view of cell-type-specific transcriptional changes in the LBDs. This combined approach is used because, while single-nucleus RNA-sequencing can address confounding by cellular composition, providing previously unattainable insight into cell-type-specific transcriptomic pathology [31,32], compared with bulk-tissue RNA-sequencing it has little ability to resolve transcriptomic diversity via splicing. This limitation arises due to the trade-off that exists between choosing a single-nucleus RNA-sequencing protocol that has high throughput but only sequences 3’ ends of transcripts versus a protocol whose library construction permits sequencing full-length transcripts but has reduced throughput [33]. Using this combined sequencing approach, we found transcriptional changes in multiple cortical cell types across the LBDs, with more differentially expressed genes and pathways identified in PDD and DLB than in PD. We also observed widespread alternative splicing, particularly in PDD and DLB, with evidence suggesting that specific splicing factors play a role in orchestrating the disease-related splicing changes. Collectively, these results identify common and distinct molecular pathology in the LBDs across several cell types and provide insight into the extent to which the LBDs represent discrete diseases with unique pathogenic processes.

## Results

### Paired single-nucleus and bulk-tissue RNA sequencing of anterior cingulate cortex in individuals with Lewy body disease

We applied single-nucleus and bulk-tissue RNA-sequencing to adjacent anterior cingulate cortex tissue sections from 28 individuals, including non-neurological control individuals and individuals with Lewy body disease (**Figure 1**). The latter were split into three disease groups, consisting of Parkinson’s disease without cognitive impairment (PD), Parkinson’s disease with dementia (PDD) and dementia with Lewy bodies (DLB), based on clinical assessments of retrospectively-reviewed case records (n = 7 per group). We sampled from the anterior cingulate cortex, as it is one of the first cortical areas to be affected by α-synuclein pathology [34,35] and a region where Lewy body densities correlate with cognitive impairment in PD [36]. Although selected individuals were matched, where possible, for demographic and pathologic factors, there were significant differences in the proportions of sexes between the groups in keeping with previous literature describing a male bias in DLB [37] (proportion female: control = 1/7, PD = 5/7, PDD = 2/7, DLB = 0/7; p-value = 0.020; Chi-squared test; **Supplementary Figure 1**, **Supplementary Table 1**). Disease duration also differed significantly between groups, with DLB cases having a shorter duration of disease before death, reflecting the fact that PDD cases have PD motor symptoms for several years before development of dementia (median disease duration in years: PD = 12, PDD = 11, DLB = 6; p-value = 0.0099; Kruskal-Wallis rank sum test; **Supplementary Figure 1**, **Supplementary Table 1**). Using this sample set, we report a total of 205,948 droplet-based single-nucleus and 24 bulk-tissue transcriptomic profiles, with an average of 1,398 genes per nucleus and 27,802 genes per bulk-tissue sample detected, respectively (Error! Reference source not found., **Supplementary Figure 3**, **Supplementary Table 1**).

**Figure 1.**
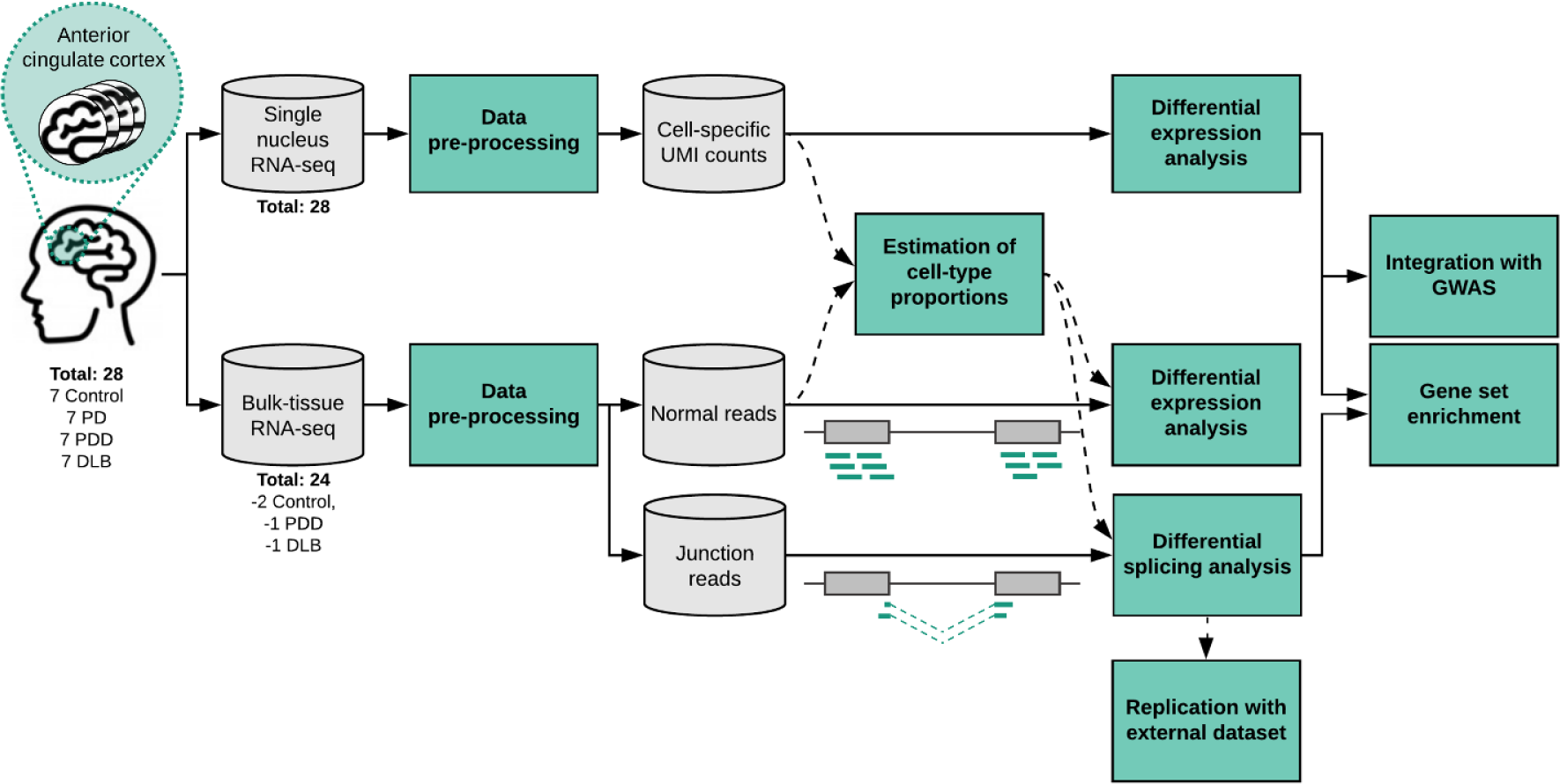
Overview of approach. In this study, anterior cingulate cortex was sampled from a cohort of 28 individuals divided equally between four groups: non-neurological controls; Parkinson’s disease without cognitive impairment (PD); Parkinson’s disease with dementia (PDD); and dementia with Lewy Bodies (DLB) (**Supplementary Figure 1**, **Supplementary Table 1**). For each individual, a frozen tissue block derived from the anterior cingulate was sectioned, with adjacent sections used for single-nucleus or bulk-tissue RNA-sequencing (Error! Reference source not found., **Supplementary Figure 3**, **Supplementary Table 1**). Following data pre-processing, single-nucleus RNA-sequencing data was used to generate cell-type-specific differential gene expression profiles and to deconvolute bulk-tissue RNA-sequencing data. Bulk-tissue RNA-sequencing was used in differential gene expression and splicing analyses, with cell-type proportions included as model covariates in both analyses. Results from single-nucleus RNA-sequencing and bulk-tissue RNA-sequencing were used in downstream gene set enrichment analyses to identify disease-relevant pathways. Furthermore, common risk variants for Alzheimer’s disease (AD), PD risk and PD age of onset (PD AOO) were mapped to cell-type-specific expression profiles and cell-type-specific differential expression.

### Increased proportions of microglia and vascular cells across Lewy body diseases

Quality control, clustering and classification of major cell types in the anterior cingulate cortex was first performed on nuclear RNA from each of the 28 individuals, after which we used the Conos framework to generate a joint graph of nuclei across all individuals [38]. Clusters were assigned to 7 broad cell types by significant overlap (Fisher’s exact test, p-value < 2.2 × 10^-16^) with a merged list of marker genes derived from two human single-cell datasets (Error! Reference source not found.**c**) [39,40]. In total, we identified 75,826 excitatory neurons, 26,467 inhibitory neurons, 46,662 oligodendrocytes, 25,726 astrocytes, 13,788 microglia, 12,497 oligodendrocyte-precursors (OPCs), and 4,532 vascular cells (which represented a merge of endothelial cells and pericytes), with each cell type consistently identified across all individuals in each disease group (**Figure 2a**, **Supplementary Figure 4a-b**).

**Figure 2.**
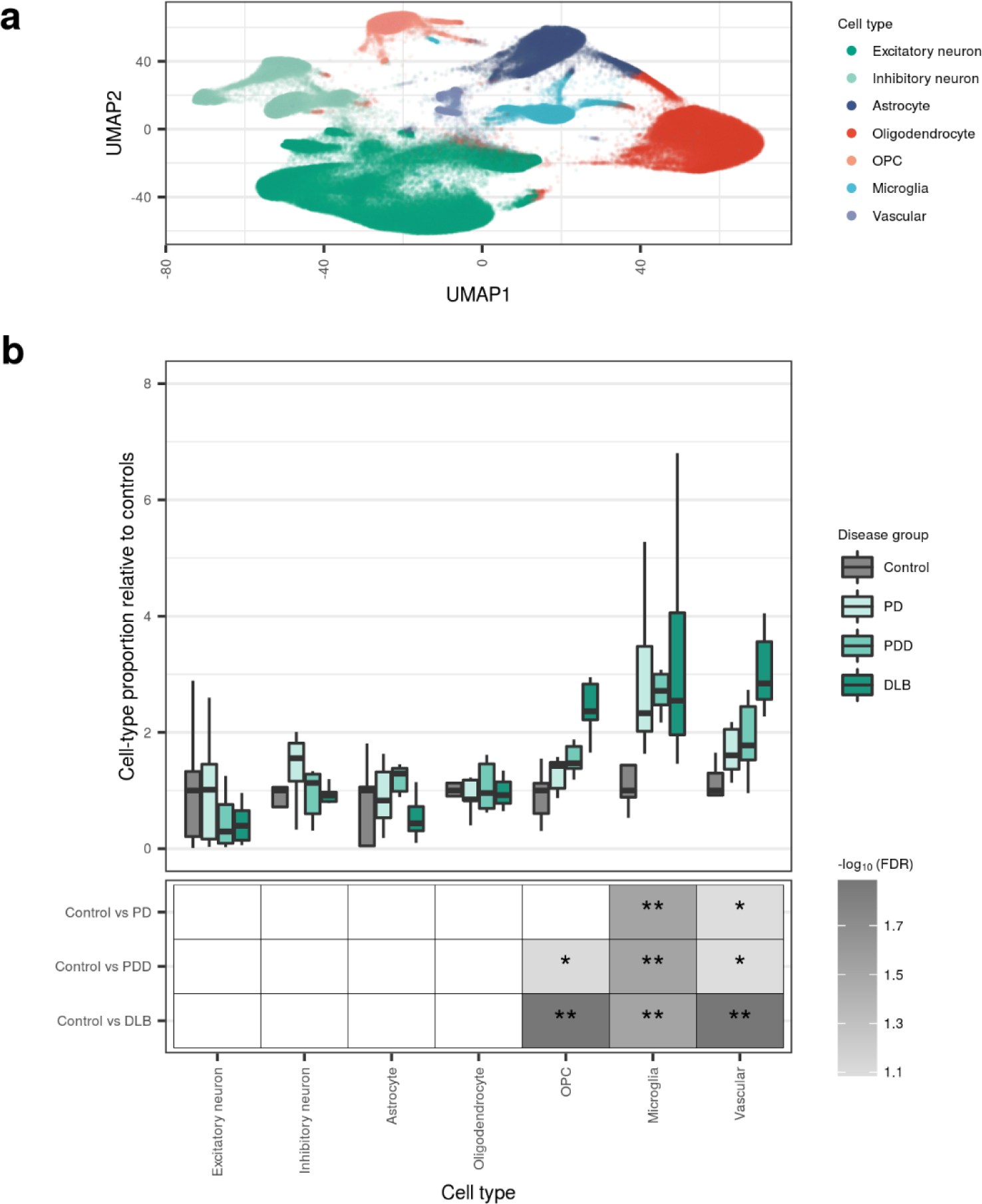
Cellular diversity of the anterior cingulate cortex across disease states. (a) Joint graph of all nuclei derived from all individuals visualised using UMAP embedding. Nuclei are coloured by cell type. **(b)** Cell-type proportions derived from Scaden deconvolution (available in **Supplementary Table 2**). Cell-type proportions (upper panel) are grouped by cell type and disease status and displayed relative to the median of controls (within a cell type). Significant differences in cell-type proportions between disease groups (lower panel) were determined using the Wilcoxon rank sum test, with FDR correction for multiple testing. Non-significant results (FDR > 0.1) were coloured white; **, FDR < 0.05; *, FDR <= 0.1.

Next, we sought to identify significant changes in the proportions of these major cell types across all disease groups. Although single-nucleus RNA-sequencing shows less sampling bias than single-cell sequencing [41], its suitability for estimation of cell-type proportions remains in question [42]. Thus, we used Scaden [43], a deep-learning-based deconvolution algorithm that can train on artificial bulk-tissue RNA-sequencing samples simulated from tissue-matched single-nucleus RNA-sequencing data, to estimate cell-type proportions across disease groups. Importantly, Scaden permitted pairing of our single-nucleus and bulk-tissue transcriptomic profiles and modelling of inter-subject variability. We observed a low overall correlation between single-nucleus-estimated and Scaden-predicted cell-type proportions (Spearman’s ρ = 0.25, p-value = 0.0009), although per-cell-type correlations were higher for some cell types (highest in microglia, Spearman’s ρ = 0.79, p-value = 8.2 × 10^-6^; Supplementary Figure 4c).

Using Scaden predictions, we identified a significantly increased proportion of microglia in all disease groups compared with the control group, and a significantly increased proportion of OPCs and vascular cells in DLB cases compared with controls (**Figure 2b**, FDR-corrected p < 0.05, Wilcoxon rank sum test). Additionally, we observed a nominally significant increase in vascular proportions in PDD and PD cases compared with controls (FDR-corrected p < 0.1, **Figure 2b**). By applying Scaden to a second, larger independent PD case-control bulk-tissue RNA-sequencing dataset [44], we were able to replicate the observed increase in microglial and vascular proportions in PD cases compared with controls (FDR-corrected p < 0.05, **Supplementary Figure 5**).

### Differential gene expression analysis highlights transcriptional alterations in multiple cell types and differentiates Lewy body dementias from PD

Differential gene expression analyses were separately performed with bulk-tissue and single-nucleus RNA-sequencing data to characterise molecular changes across the disease groups (**Materials** and methods). Following correction for changes in Scaden-predicted cell-type proportions in bulk-tissue gene expression, only 60 genes (53 unique genes) were found differentially expressed (DE) across the six pairwise comparisons (FDR < 0.05, **Supplementary Table 3**). Despite the low number of bulk-tissue DE genes identified, we noted that gene expression adjusted for cell type and experimental covariates resulted in much clearer clustering of samples by disease group (as determined through visual inspection) compared with uncorrected gene expression and gene expression adjusted for experimental covariates alone (**Supplementary Figure 6a-c**). Notably, separation of disease groups was primarily observed on the same axis of variation (i.e. the first principal component, PC1), suggesting that (i) the genes contributing most to variation between groups are similar across disease groups, and thus PD, PDD and DLB may represent a neuropathological continuum and (ii) that there are gene expression changes between disease groups that are independent of differences in cell-type proportions (**Supplementary Figure 6a-c**). Using pathway enrichment, we found that the top 100 genes contributing to PC1 were associated with immune-related GO terms (e.g. peptide antigen binding and MHC protein complex), as well as terms relating to endocytic vesicles and unfolded protein binding (**Supplementary Figure 6d**, **Supplementary Table 4**).

Consistent with the view that gene expression changes exist between disease groups independent of differences in cell-type proportions, using single-nucleus RNA-sequencing data, 9,242 unique genes were found DE across cell-type-specific pairwise comparisons (all six pairwise comparisons,|log_2_(fold change)| > log_2_(1.5), FDR < 0.05, **Supplementary Table 5**). Focusing only on comparisons with the control group, these analyses highlighted three main themes.

First, differential gene expression was widespread and involved glia and neurons. While we found that DE genes were detected across all three case-control comparisons and across all major cell types, the largest numbers of DE genes were observed in excitatory neurons, followed by oligodendrocytes (**Figure 3a**). In fact, across case-control comparisons, the number of DE genes identified in oligodendrocytes exceeded that in inhibitory neurons by a factor of up to 11.4-fold (depending on the case-control comparison; **Figure 3a**). Comparison of the Lewy body diseases to each other yielded similar results; that is, transcriptional alterations across all major cell types, but with the largest number of DE genes observed in excitatory neurons, followed by oligodendrocytes (**Supplementary Figure 7**).

**Figure 3.**
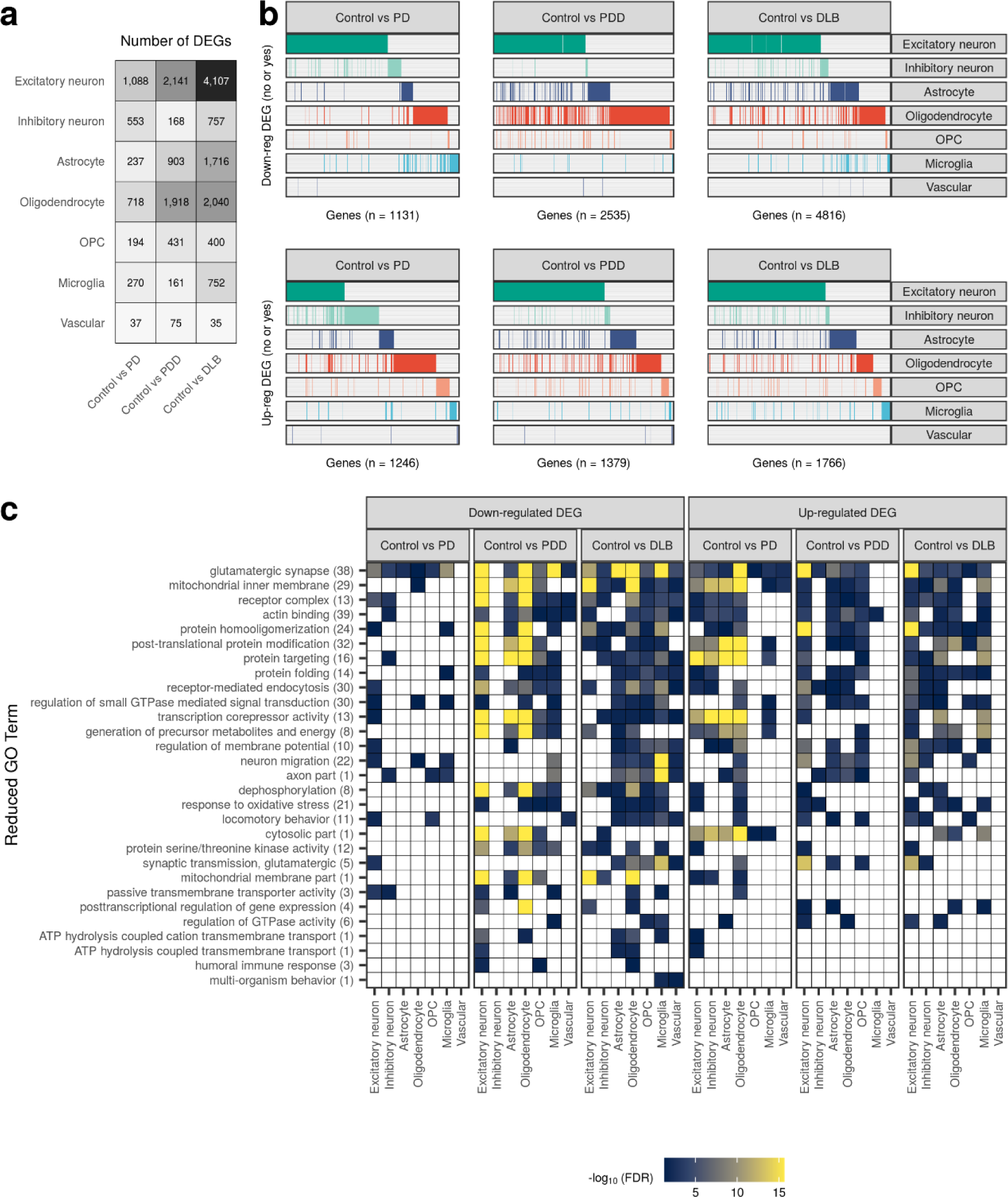
Cell-type-specific gene expression changes and pathway enrichments across disease states. (a) Number of differentially expressed (DE) genes across each cell type in pairwise comparisons of disease groups to the control group (|log_2_(fold change)| > log_2_(1.5), FDR < 0.05). The intensity of the grey colour is proportional to the number of DE genes. **(b)** Binary plot indicating with bars whether a gene (column) is down-regulated (upper panel) or up-regulated (lower panel) in a given cell type (rows). Number of DE genes in each comparison indicated on the x-axis. **(c)** Reduced GO terms associated with cell-type-specific down- and up-regulated DE genes identified across pairwise comparisons of disease groups with the control group. Due to the magnitude of pathway enrichments, original GO term enrichments (referred to as “child terms”) were grouped using semantic similarity. The number of enriched child GO terms assigned to each reduced parent term across all cell types and comparisons in the panel is indicated in parentheses on the y-axis. Reduced GO terms were ordered on the y-axis by the number of cell types and comparisons in which the term was found enriched. The fill of each tile indicates the -log10(FDR) of the most significant child term associated with the parent term within that comparison/cell type. Non-significant results (FDR > 0.05) were coloured white. Results for pairwise comparisons between disease groups are displayed in **Supplementary Figure 7**. All cell-type-specific DE genes and pathway enrichments are available in **Supplementary Table 5** and **Supplementary Table 6**, respectively.

Second, DE genes were commonly specific to a cell type. Indeed, of the 1,131, 2,535 and 4,816 down-regulated DE genes identified across comparisons of PD, PDD and DLB with control, 79%, 66% and 67%, respectively, were DE in only one cell type (**Figure 3b**). Among up-regulated DE genes, these percentages ranged from 74-76% across the three case-control comparisons.

Third, the Lewy body dementias, as distinct from PD, were characterised by the predominant down-regulation of gene expression relative to control in most cell types; the only exception were inhibitory neurons in PDD, where the number of up-regulated DE genes exceeded the number of down-regulated DE genes (**Figure 3a-b**). Furthermore, the transcriptomic profile of the two Lewy body dementias was very similar, with 303 down-regulated and 87 up-regulated DE genes identified in a comparison of DLB with PDD (**Supplementary Figure 7**). In contrast, comparisons of the two Lewy body dementias with PD identified > 2,000 down-regulated and > 1,000 up-regulated DE genes, suggesting that while there are transcriptional commonalities between PDD and DLB, PD is transcriptionally distinct from the Lewy body dementias in the anterior cingulate cortex.

Pathway enrichment was used to explore the biological implications of cell-type-specific differential gene expression. Focusing on case-control comparisons, we found that down- and up-regulated DE gene sets were enriched for 306 and 272 GO terms, respectively (each pathway was only counted once, even if it appeared across > 1 case-control comparison). Using measures of semantic similarity to cluster GO terms, and thus reduce pathway redundancy, we identified 29 down-regulated and 27 up-regulated GO terms (**Figure 3c**, **Supplementary Table 6**). Despite the high proportion of cell-type-specific DE genes, we identified GO terms that were perturbed across multiple cell types in a given case-control comparison. For example, in comparisons of PD with control, terms related to glutamatergic synapses, the mitochondrial inner membrane, and post-translational protein modification were enriched across ≥ 5 cell types . These commonalities in GO term enrichment were a feature of both down- and up-regulated DE gene sets but were more apparent among (i) down-regulated DE gene sets and (ii) comparisons of PDD and DLB with control, with pathway perturbations affecting a median of 3-5 cell types, as compared with 1-3 in comparisons of PD with control (**Supplementary Figure 8a**). Furthermore, we noted that consistent with the high number of DE genes detected for excitatory neurons, a high number of enriched pathways were observed in this cell type across all case-control comparisons, particularly in PDD and DLB (**Supplementary Figure 8b**). This observation was even more pronounced in comparisons of the Lewy body dementias with PD, where the number of enriched pathways identified in excitatory neurons was almost 2-fold higher than the second most-affected cell type. Overall, this analysis served to highlight disproportionately large transcriptional differences in PDD and DLB, as compared with PD, particularly in excitatory neurons and, to a lesser extent, oligodendrocytes.

### Genes and pathways genetically associated with PD implicate physiological variability of SNCA expression in selective vulnerability of neurons

Many of the GO terms enriched among down- and up-regulated genes, such as receptor-mediated endocytosis, have been previously implicated in PD. With this in mind, we narrowed our focus to the cell-type specific expression of genes and pathways genetically associated with PD pathogenesis [45,46].

PD-associated genes were derived from a recent review of mutations that have been reported to cause PD, including well-known examples such as *SNCA* [45]. Of the 21 genes considered, 13 were DE in at least one major cell type and one case-control comparison (**Figure 4a**). For example, excitatory neurons, inhibitory neurons, astrocytes and oligodendrocytes all showed significant up-regulation of *SNCA* in PD cases when compared with controls (fold change: 0.64 – 1.30; FDR: 2.6 × 10^-7^ – 7.2 × 10^-157^, **Figure 4a**).

**Figure 4.**
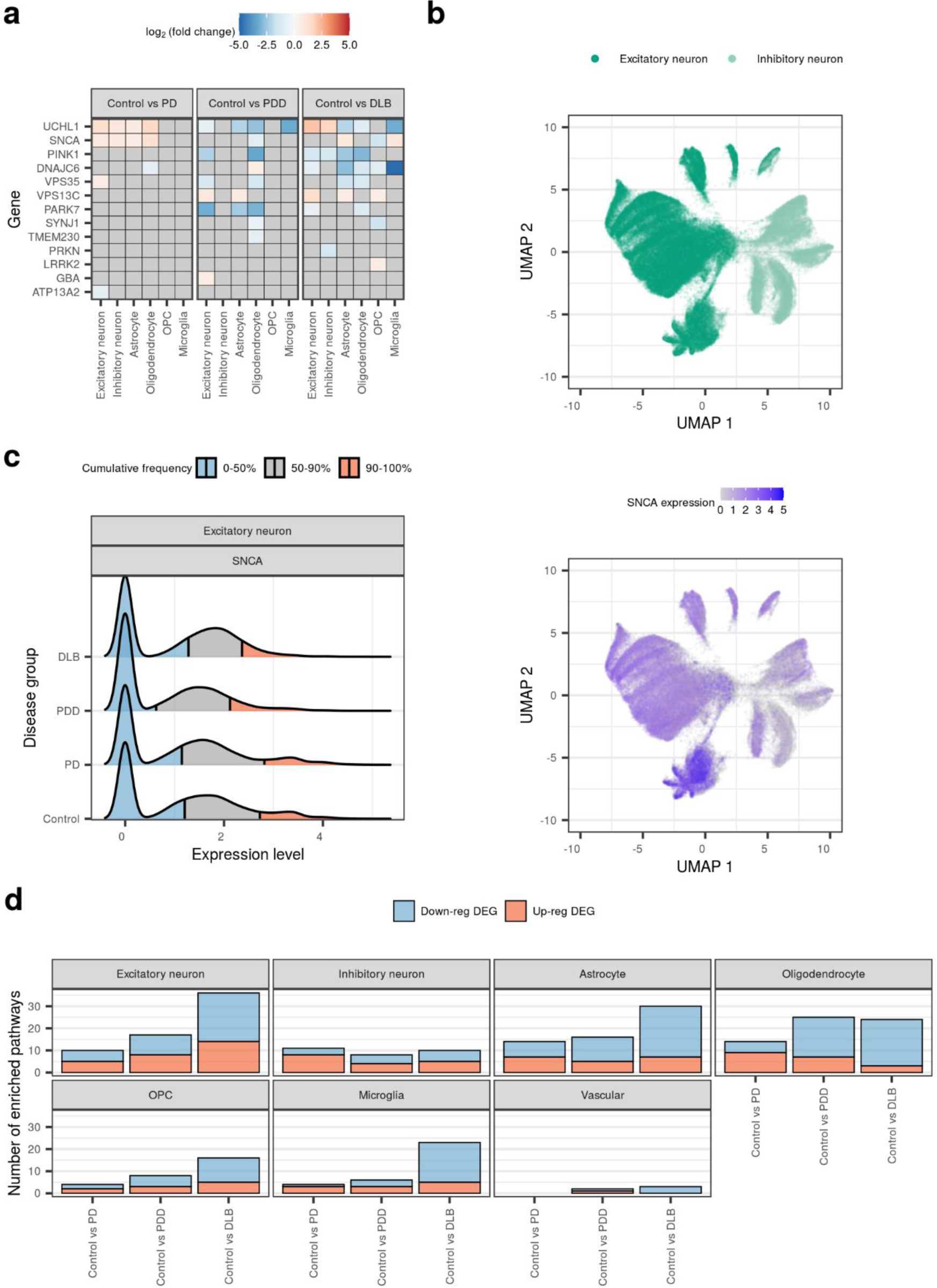
Cell-type-specific alterations of PD-associated genes and pathways. (a) Differential expression of PD-associated genes (associated by mutations reported to cause PD [45]) across cell types and pairwise comparisons of disease groups with the control group. Fill of the tile indicates the log_2_(fold change), with non-significant results (FDR > 0.05) coloured grey. **(b)** UMAP plot of excitatory and inhibitory neurons (upper panel, 102,293 nuclei), with *SNCA* expression levels (lower panel). **(c)** Ridgeline plot of distribution of *SNCA* expression levels in excitatory neurons across disease groups. Distributions have been split into 3 cumulative quantiles, highlighting where 0-50%, 50-90% and 90-100% of the nuclei in each disease group lie. **(d)** Number of enriched pathways (FDR < 0.05) identified using cell-type-specific down- and up-regulated DE genes from each pairwise comparison together with 46 PD-associated pathways (associated in a large-scale polygenic risk score-based assessment of 2,199 gene sets [46]).

There is robust genetic evidence linking increased *SNCA* dosage to PD pathogenesis, including (i) duplication and triplication events in the *SNCA* gene that underlie autosomal dominant forms of PD [47,48] and (ii) the association of PD risk loci with increased *SNCA* expression [49,50]. In view of this evidence, we further explored *SNCA* expression, finding that, while *SNCA* expression was up-regulated in PD in all four cell types with a similar fold change (**Figure 4a**), *SNCA* expression in control individuals was highly variable across cell types (**Supplementary Figure 9**). This variability in control *SNCA* expression extended to (i) the proportion of nuclei expressing *SNCA*, with 61% of excitatory neurons expressing *SNCA*, as compared with < 22% across all other cell types, and (ii) the range of observed *SNCA* expression, which was wider in excitatory neurons compared with all other cell types (**Supplementary Figure 9**). These differences in cell-type-specific *SNCA* expression were particularly apparent between inhibitory and excitatory neurons, irrespective of disease group, with a higher proportion of excitatory neurons expressing *SNCA* (**Figure 4b**, **Supplementary Figure 9**). Furthermore, these differences were visible in a cell type across disease groups. Indeed, *SNCA* expression in excitatory neurons from the Lewy body dementias, as compared with the control group, was marked by (i) a decrease in the proportion of *SNCA*-expressing nuclei in PDD and (ii) a shift in the expression range of the top 10% highest-expressing nuclei to lower levels of *SNCA* expression (**Figure 4c**). This was not, however, the case for PD, which maintained a similar distribution of *SNCA* expression to the control group, with a slight shift in the expression range of the top 10% highest-expressing nuclei to higher levels of *SNCA* expression. The absence of a population of cells expressing higher levels of *SNCA* suggests that variability in *SNCA* expression within control ranges may contribute to the selective vulnerability of subpopulations of excitatory neurons to Lewy body pathology.

PD-associated pathways were leveraged from a recent study identifying 46 pathways implicated in PD through pathway-specific polygenic risk score and rare variant burden analyses [46]. Based on case-control comparisons, we found that pathways that have been genetically associated with PD causation (such as terms related to synaptic transmission and vesicle-mediated transport) were dysregulated in all major cell types, with the exception of vascular cells, wherein only 3 pathways were implicated (**Figure 4d**, **Supplementary Figure 10**, **Supplementary Table 7**). We noted that the number of dysregulated pathways tended to increase with increasing clinical disease severity (i.e. PD < PDD < DLB) in excitatory neurons and glia, but not inhibitory neurons and vascular cells, supporting the notion of a disease spectrum. In general, fewer pathways were dysregulated in inhibitory neurons, with 12/46 pathways dysregulated in at least one case-control comparison, as compared with excitatory neurons, astrocytes and oligodendrocytes (23-27/46 pathways).

### Differentially expressed genes in glia enrich for heritability of PD age of onset and risk

To identify cell types through which common genetic variants associated with PD risk and dementia may be acting, we used Hi-C-coupled Multi-marker Analysis of GenoMic Annotation (H-MAGMA) [51] and stratified LD score regression (sLDSC) [52]. As age of PD onset is correlated with clinical progression [53–55], and there is a significant negative genetic correlation between the GWAS for PD age of onset (AOO) and PD risk [56], we included both GWASs in our analysis. Further, given the potential cooccurrence of Alzheimer’s disease (AD) pathology in the Lewy body dementias, we used a recent late-onset AD GWAS [57].

Genetic association analyses with H-MAGMA and sLDSC were run with two sets of annotations: (i) the top 10% most cell-type-specific genes from each disease group and (ii) cell-type-specific DE genes (|log_2_(fold change)| > log_2_(1.5), FDR < 0.05). The latter were tested on the basis that DE genes better capture gene expression signatures representative of a given disease state. Using the top 10% most cell-type-specific genes, we observed a significant association between AD genetic risk and genes highly expressed in microglia derived from control, PD and PDD groups (control, FDR_LDSC_ = 0.038; PD, FDR_LDSC_ = 0.019; PDD, FDR_LDSC_ = 0.035; **Figure 5a**; **Supplementary Table 8**), replicating previous literature [57–59]. Furthermore, we observed a significant association between genetic determinants of PD age of onset and genes highly expressed in OPCs derived from the DLB group (FDR_HMAGMA_ = 0.022) and PD genetic risk and genes highly expressed in oligodendrocytes (a cell type of increasing interest to the PD field [58,59]) derived from the control group (FDR_HMAGMA_ = 0.013).

**Figure 5.**
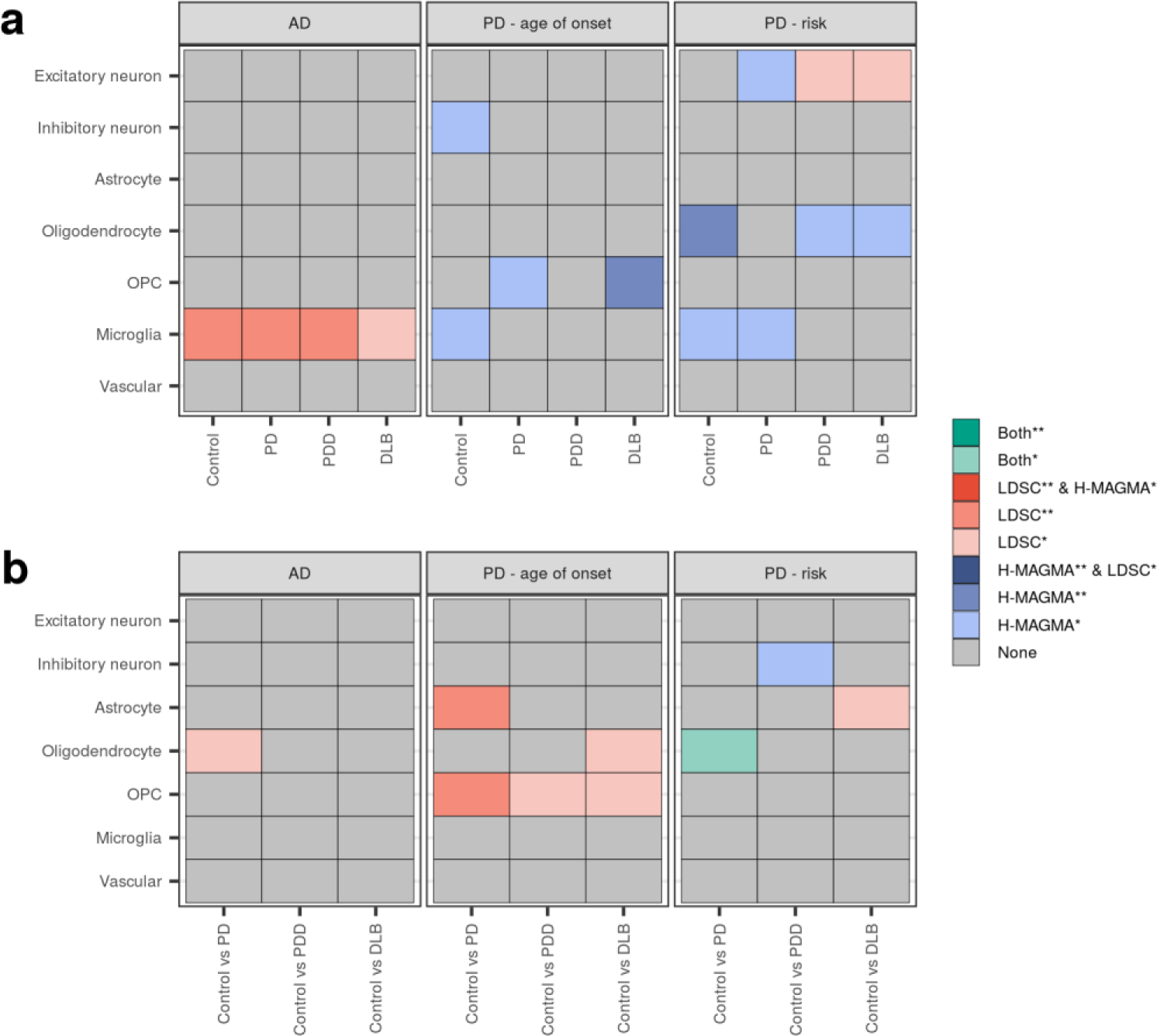
Genetic associations with top 10% most cell-type-specific genes and cell-type-specific differentially expressed genes. Genetic associations using **(a)** the top 10% most cell-type-specific genes in each disease group and **(b)** cell-type-specific differentially expressed genes in disease comparisons with controls. Two methods were used to identify associations: Hi-C-coupled MAGMA (H-MAGMA) and stratified LD score regression (sLDSC). The heatmap is coloured by degree of significance with both or either method, with * and ** indicating nominal significance (unadjusted p-value < 0.05) or significance (FDR-corrected p-value < 0.05; corrected for number of cell types tested). Results available in **Supplementary Table 8**. AD, Alzheimer’s disease; PD, Parkinson’s disease.

Using cell-type-specific DE genes, we identified a significant association between genetic determinants of PD age of onset and genes found DE in astrocytes and OPCs from comparisons of PD with control (astrocytes, FDR_LDSC_ = 0.0085; OPCs, FDR_LDSC_ = 0.0085; **Figure 5b**). Splitting differentially expressed genes by their direction of effect showed that this signal was driven by up-regulated genes (**Supplementary Figure 11**). In addition, we identified a nominal association using both methods between PD genetic risk and genes found DE in oligodendrocytes from comparisons of PD with control (P_HMAGMA_ = 0.011, P_LDSC_ = 0.041; **Figure 5b**), which was driven by up-regulated genes (FDR_HMAGMA_ = 0.013, P_LDSC_ = 0.044; **Supplementary Figure 11**). Finally, we noted that genes up-regulated in excitatory neurons from comparisons of PDD with control were significantly associated with PD genetic risk (FDR_LDSC_ = 0.040; **Supplementary Figure 11**).

### Differential splicing distinguishes PDD from DLB and highlights the role of specific RNA-binding proteins

Given the limitations of single-nucleus RNA-sequencing in the detection of splicing, we applied Leafcutter to our bulk-tissue RNA-sequencing to assess differential splicing (DS) [60]. Leafcutter captures changes in local splicing events through construction of intron clusters, wherein overlapping introns are connected by the splice junction(s) they share. We identified a total of 4,656 DS intron clusters in 3,751 genes (FDR < 0.05, |ΔPSI| ≥ 0.1; **Supplementary Table 9**) across all pairwise comparisons, with the highest number identified in comparisons of DLB with control or PD (**Supplementary Figure 12a**). Notably, between 28-32% of DS events were partially annotated with respect to the reference transcriptome, with splicing events including novel donor or acceptor splice sites, novel exon skip and novel combination events (**Supplementary Figure 13a-b**). We were, however, able to detect these events in larger control cohorts suggesting that they represent biologically relevant splicing (**Supplementary Note**, **Supplementary Figure 13c-d**).

DS genes showed a significant enrichment in oligodendrocytes across comparisons of all disease groups with the control group (i.e. these genes had higher expression in oligodendrocytes than expected by chance), an observation that we replicated using the same external PD case-control bulk-tissue RNA-sequencing dataset used in replication of deconvolution results (**Figure 6a**, **Supplementary Note**, **Supplementary Figure 15a**, **Supplementary Table 10**). In contrast, enrichments in other cell types appeared to be disease specific (**Figure 6a**). For example, only genes found DS in comparisons of PD with control or DLB with PD enriched in astrocytes. Notably, as the only pairwise comparison, DS genes from DLB compared with PDD consistently enriched in all excitatory neuron annotations. Pathway enrichments were observed across 4/6 pairwise comparisons (no enrichments were observed in comparisons of PD or PDD with control; **Supplementary Figure 12b**, **Supplementary Table 11**). Pathways that were shared across comparisons of DLB with control, PD and PDD, included terms related to endosomes and enzyme activity (in particular, GTPase activity), mirroring terms highlighted both by replication analyses and by pathway analysis of single-nucleus DE genes (**Figure 6b**, **Supplementary Note**, **Supplementary Figure 12b**, **Supplementary Figure 15b**).

**Figure 6.**
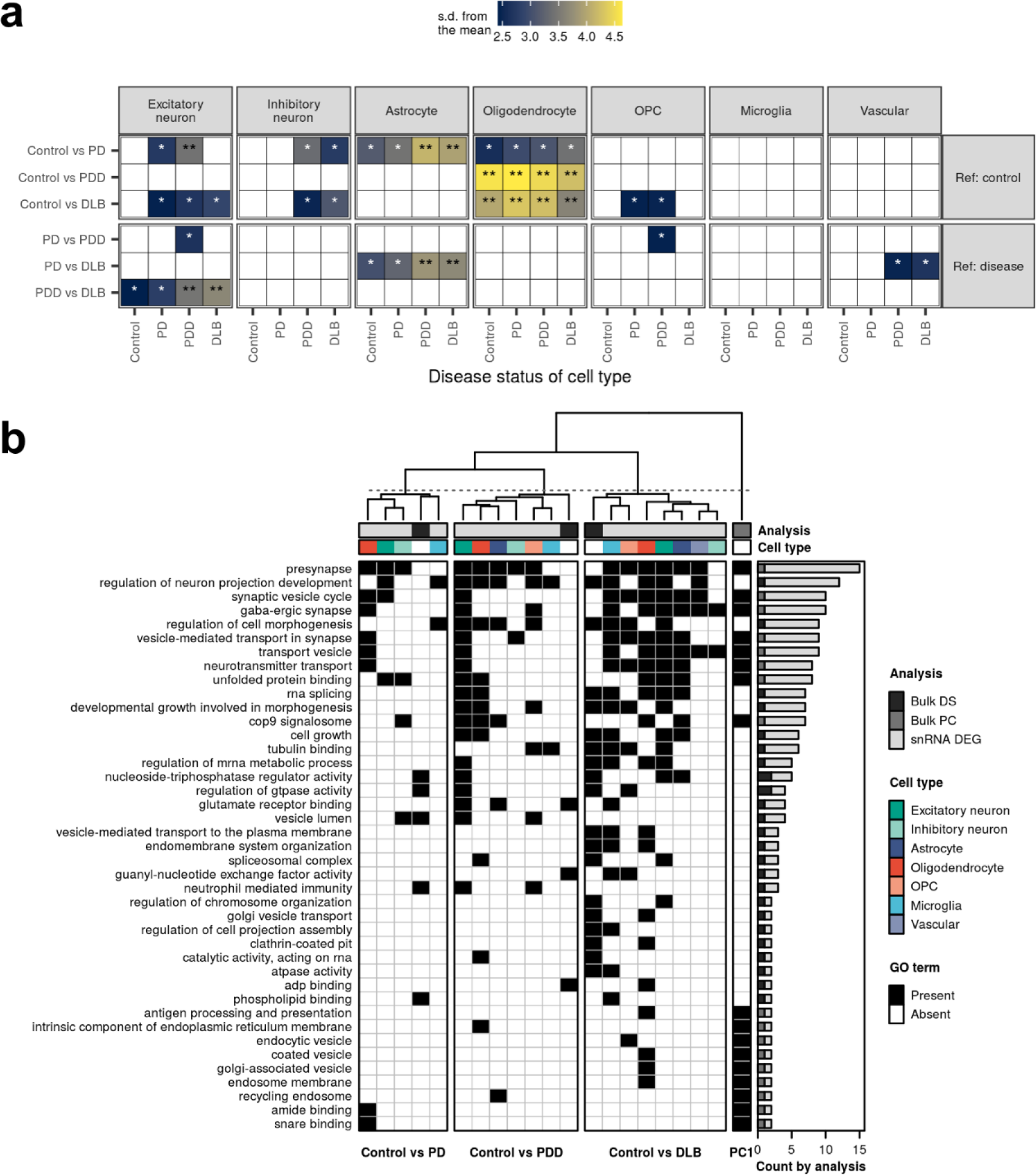
Cell-type enrichments of differentially spliced genes and pathway sharing across analyses. (a) Enrichment of the top 100 DS genes (FDR < 0.05, |ΔPSI| >= 0.1, with rank determined by |ΔPSI|) in cell types derived from each disease group. Enrichments were determined using EWCE. The x-axis denotes the disease status of the cell type in question, while the y-axis denotes the groups compared in the differential splicing analysis. Pairwise comparisons have been grouped by whether diseased individuals are compared with control individuals (Ref: control) or other diseased individuals (Ref: disease). Tiles were coloured by standard deviations from the mean, which indicate the distance (in standard deviations) of the target list from the mean of the bootstrapped samples. Multiple test correction was performed across EWCE results using FDR. Non-significant results (FDR > 0.05) were coloured white. ***, FDR < 0.001; **, FDR < 0.01; *, FDR < 0.05. All results available in **Supplementary Table 10**. **(b)** Clustering of shared pathway enrichments using genes identified across the three main analyses (represented by grey bar entitled, “Analysis”). These included: bulk-tissue differential splicing (“Bulk DS”, **Supplementary Figure 12**); gene contributions to bulk-tissue gene expression PC1 (“Bulk PC”, **Supplementary Figure 6**); and single-nucleus differential expression (“snRNA DEG”, Figure 3). Pathways (in rows) from all three analyses were filtered to include only those that appear across more than one type of analysis. Pathways are ordered from highest to lowest by the number of gene sets in which they are enriched (as displayed in the bar plot on the right-hand side). Gene sets (in columns) are clustered using hierarchical clustering on the Pearson correlation between gene sets (pathways were encoded with a binary 1 for “Present” or 0 for “Absent”, represented on the plot by black and white, respectively). Gene sets derived from differential splicing (Bulk DS) were collapsed across our own dataset and the validation dataset, resulting in one gene set (column) per pairwise comparison. Likewise, gene sets derived from up- and down-regulated single-nucleus DE gene sets were collapsed across cell types (represented by the coloured bar entitled, “Cell type”), such that each cell type was represented by a single column. Pathway overlaps using pairwise comparisons between disease groups are displayed in **Supplementary Figure 16**.

Visualisation of pathway sharing across gene sets derived from the three analyses (bulk-tissue differential splicing, gene contributions to bulk-tissue gene expression PC1 and single-nucleus differential expression) demonstrated limited sharing between the two bulk-tissue analyses (the exceptions being “presynapse”, “transport vesicle”, “coated vesicle”, and “endosome membrane”; **Figure 6b**; **Supplementary Figure 16**). Notably, pathway analysis of DS genes from DLB compared with PDD implicated a much wider breadth of pathways compared with pathway analysis of single-nucleus DE genes from the same comparison, and indeed, no pathways overlapped between the two analyses in this pairwise comparison (**Supplementary Figure 16**). This observation suggests that differences between PDD and DLB are not sufficiently captured by consideration of gene expression alone.

Patterns of pathway sharing between each of the bulk-tissue analyses and single-nucleus differential expression highlighted highly shared terms related to synaptic function, unfolded protein binding, and vesicle transport. Of note, RNA splicing was (i) jointly implicated by differential splicing and single-nucleus differential expression derived from excitatory neurons, oligodendrocytes, astrocytes and microglia in comparisons of DLB with control and (ii) separately implicated by single-nucleus differential expression derived from excitatory neurons and oligodendrocytes in comparisons of PDD with control (**Figure 6b**). Together with the abundant differential splicing observed, these results indicated that dysregulation of splicing factors may play a role in the pathogenesis of LBDs.

To further investigate this observation, we used a catalogue of known RNA-binding protein (RBP) binding motifs from the ATtRACT database [61], and defined introns by their proximal intronic regions (the 50 nt of an exon and 500 nt of an intron flanking the 5’ and 3’ splice sites), which are an important region for splicing regulation [62]. Proximal intronic regions from DS introns were compared with non-DS introns across each pairwise comparison, identifying a total of 4 RBP binding motifs with a significant enrichment in DS proximal intronic regions from at least one pairwise comparison (**Supplementary Table 12**). Among these was the consensus sequence GGGGGGG in DS proximal intronic regions from PDD comparisons with control (Bonferroni-adjusted p-value = 0.000601; **Supplementary Table 12**). This sequence is targeted by 17 RBPs from the ATtRACT database (including several members of the hnRNP family, such as *HNRNPC* and *FUS*), as well as RBPs not included in the database, such as *RBM25* [63,64]. Notably, *RBM25* was found DS across comparisons of PDD with control in our own dataset and the validation dataset (in-house, clu_26788, FDR-adjusted p-value = 0.00653; SRP058181, clu_12260, FDR-adjusted p-value = 0.0499; **Supplementary Table 9**). Furthermore, the consensus sequence GAAGGAA, targeted by *HNRNPM*, was enriched in DS proximal intronic regions from comparisons of DLB with control and PD (Bonferroni-adjusted p-values, control vs DLB = 0.0141, PD vs DLB = 0.00133). Finally, two consensus sequences, CUGGAUU and CUAACCCUAA targeted by *SRSF9* and *PCBP2*, respectively, were enriched in DS proximal intronic regions from comparisons of DLB with PDD (Bonferroni-adjusted p-values, CUGGAUU = 0.000958, CUAACCCUAA = 0.0174). Of note, SRp30c (encoded by *SRSF9*) has been shown to interact with hTRA2-β (encoded by *TRA2B*) [65,66], which targets the consensus sequence AAGAAGAAGAA, which we also found to be nominally enriched in DS proximal intronic regions from comparisons of DLB with PDD (Bonferroni-adjusted p-value = 0.0865).

Overall, these results highlighted (i) the abundant levels of alternative splicing, particularly in PDD and DLB, with evidence to suggest that certain splicing factors may play a role in orchestrating these disease-related splicing changes and (ii) that differential splicing, particularly in comparisons of DLB with PDD, captures additional features of disease-related perturbations, which were not captured by single-nucleus differential gene expression.

## Discussion

Here, we applied paired bulk-tissue and single-nucleus RNA-sequencing to transcriptomically profile PD, PDD and DLB. Using this approach, we (i) found transcriptional differences relative to controls for multiple cell types across the LBDs, with PDD and DLB more severely affected than PD; (ii) observed high levels of alternative splicing, particularly in PDD and DLB; and (iii) identified splicing factors, with links to other dementia-related neurodegenerative diseases, that may coordinate these disease-related splicing changes. Together, these results highlight transcriptomic commonalities and distinctions between the LBDs, which can be used to inform our understanding of the relationship between these three clinical disorders.

Existing transcriptomic studies of the LBDs have relied on bulk-tissue analyses and profiled each disease separately, limiting our understanding of the molecular landscape of these diseases individually and in relation to one another. In addition, few initiatives have addressed genome-wide assessment of splicing in this context, despite studies implicating alternative splicing as a disease mechanism in monogenic and sporadic forms of PD [50,67], and complex disease, in general [68]. Using multiple sequencing and analytic approaches, our analyses had the potential to identify differences between the LBDs attributable to changes in cell-type proportions, cell-type-specific gene expression and bulk-tissue splicing. While we found that increases in microglial and vascular cell-type proportions were a feature of LBDs, with the microglial increase mirroring results from an RNA-sequencing-based study of PD modelling cellular composition [69], these increases did not distinguish among the LBDs. In contrast, cell-type-specific differential gene expression and bulk-tissue differential splicing distinguished PD from the Lewy body dementias, with PDD and DLB demonstrating a higher degree of commonality. These results suggest that irrespective of when dementia onset occurs in the disease process it gives rise to similar end-stage, post-mortem transcriptomic signatures in the anterior cingulate cortex.

It is notable that bulk-tissue differential splicing (i) was a prominent feature of the LBDs, (ii) discriminated between PD and the Lewy body dementias and (iii) provided evidence of relationships with other neurodegenerative diseases clinically associated with dementia. Enrichment analyses using DS genes associated with each of the three LBDs revealed shared cell-type associations, such as the differential splicing of genes highly expressed in oligodendrocytes, as well as disease-specific cell-type and pathway associations. Indeed, splicing analyses highlighted pathways relating to GTPase activity and regulation across several pairwise comparisons involving DLB, perhaps due to their role in a range of cellular processes that have been implicated in PD, such as clearance of Golgi-derived vesicles through the autophagy-lysosome system, mitochondrial fission and fusion, and p38 MAPK signalling [46,70]. RNA splicing was additionally associated with the Lewy body dementias, by both differential splicing and single-nucleus differential expression. To further investigate these observations, we assessed RBP binding motif enrichment to identify potential upstream regulators of splicing. All four significantly enriched RBP binding motifs were targeted by RBPs that have been implicated to varying degrees in neurodegenerative diseases, with *HNRNPC* implicated in AD [71], and *FUS, HNRNPC, HNRNPM* and *PCBP2* associated with frontotemporal dementia (FTD) [72]. Furthermore, not only has *PCBP2* (encoding hnRNP E2) been found to TDP-43 pathology in specific pathological subtypes of FTD [73], but *SRSF9* together with *TRA2B* are implicated in tau splicing [66]. Given that both Lewy body dementias are characterised by co-pathology [20,21], including tau and TDP-43 pathology, we speculate whether dysregulation of splicing might be one of the drivers of this co-pathology. Further studies will be required to understand whether this is the case.

Looking at cell-type-specific differential gene expression, the most prominent difference between the LBDs was the widespread down-regulation of genes and pathways in the Lewy body dementias, as compared with PD. In genetic association analyses, these genes did not enrich for genetic determinants of PD age of onset or PD risk, suggesting that this down-regulation is a consequence of the disease process, as opposed to a cause. In contrast, up-regulated genes (identified primarily in comparisons of PD with control) enriched for genetic determinants of PD age of onset and PD risk, highlighting known (OPCs/oligodendrocytes [58,59]) and new (astrocytes) cell types in PD pathogenesis. In fact, common to all three LBDs was the presence of transcriptional alterations across multiple cell types. While DE genes were found to be largely cell-type-specific (i.e. DE in only one cell type), these genes converged on similar pathways, with GO terms found to be perturbed across multiple cell types in a given case-control comparison. Restricting to genes and pathways genetically associated with PD (which arguably are more likely to be causal), we similarly saw multiple cell-type involvement across all three LBDs, albeit with some suggestion of a hierarchy of increasing perturbation in excitatory neurons and glia (i.e. PD < PDD < DLB). Together, these results suggest the involvement of multiple cell types in LBD pathogenesis, and potentially indicate a common regulatory response across cell types in each disease.

While we observed transcriptional alterations in multiple cell types, some cell types, such as excitatory neurons and oligodendrocytes, were more strongly impacted than others (most notably, inhibitory neurons), implying some degree of selective vulnerability. In support of this observation, expression of *SNCA* (encoding α-synuclein, the major component of Lewy bodies [74]) in excitatory neurons from the Lewy body dementias, as compared with the control group, was marked by a decrease in the proportion of *SNCA*-expressing nuclei in PDD and a shift in the expression range of the top 10% highest-expressing nuclei to lower values. While we recognise that this is an observational study, it is tempting to speculate that (i) variability in physiological levels of *SNCA* may impact on pathogenesis, an area of research that has received far less attention as compared with increased *SNCA* dosage [47–50], and (ii) that the absence of cells expressing high physiological levels of *SNCA* may contribute to the selective vulnerability of subpopulations of excitatory neurons to Lewy body pathology.

There are several limitations to this work that emphasise key areas for future work; the most important are the study of one brain region in diseases that gradually affect multiple brain regions and the small size of the cohort used. Where possible, we attempted to validate results in larger independent control and case-control studies, but larger studies covering more brain regions will be needed in the continuing assessment of the LBDs.

Among technological limitations, a known issue in single-nucleus RNA-sequencing is the depletion of transcripts that preferentially enrich in the cytoplasmic compartment, such as transcripts that localise to neuronal dendrites [41] and signatures of microglial activation [75]. This limitation has implications both for differential gene expression, but also downstream deconvolution and indeed, the use of single-nucleus RNA-sequencing as a reference was found to decrease the performance of three deconvolution algorithms (including Scaden) on post-mortem human brain data [43]. This limitation stresses the importance of relating cell types defined by single-nucleus RNA-sequencing back to their spatial phenotypes, a process for which the emerging field of spatial transcriptomics will be instrumental in resolving [76]. Our results provide clear hypotheses to test using spatial transcriptomics both for cell-type-specific DE analysis and analysis of differential cell-type proportions.

Among methodological limitations, we recognise that RBP binding motif enrichment oversimplifies the biology of RBPs. A common feature of RBPs is the presence of multiple RNA-binding domains, which are thought to interact with repeating motifs spaced apart on pre-mRNA transcripts [64,77]; this feature is not captured in the current analysis. Similarly, our analyses do not account for sequence context [64] (e.g. flanking nucleotide composition, repeated motifs, RNA structure) and thus cannot distinguish between RBPs that bind similar motifs. Developing tools that could address this *in silico* represents an opportunity to identify additional regulators of splicing in the LBDs.

In summary, our comprehensive transcriptomic analysis of all three LBDs highlights the complex, multi-cell-type transcriptional response to Lewy body pathology and LBD co-pathologies. Furthermore, it identifies post-mortem molecular signatures in the anterior cingulate cortex that distinguish PD from the two Lewy body dementias, such as perturbation of RNA splicing, a mechanism linked to several dementia-related neurodegenerative diseases. Together, these findings have important implications for the design of RNA-targeted therapies for these diseases and highlight a potential molecular “window” of therapeutic opportunity between the initial onset of PD and subsequent development of Lewy body dementia.

## Materials and methods

### Sample selection

Individuals with clinical parkinsonism and/or Dementia with Lewy Bodies (DLB) and pathologically confirmed PD were obtained from the Parkinson’s UK Tissue Bank. Clinical assessment of individuals was carried out on clinical notes collated retrospectively using records from movement disorder neurologists, neurosurgeons, psychiatrists, geriatricians, PD nurse specialists and general practitioners. Clinical parkinsonism was defined using the current MDS task force criteria [78] and Lewy body dementia by the most recent clinical diagnostic criteria for PDD and DLB [15,16]. The one-year rule, alongside positive clinical features for DLB (spontaneous parkinsonism, REM-sleep behaviour disorder, fluctuating cognition and complex visual hallucinations) were used to separate individuals with PDD and DLB. Pathologic assessment was performed on representative tissue sections from recommended brain regions in the Braak α -synuclein [79] and Braak tau [80] staging systems as part of the routine diagnostic process for the Parkinson’s UK Tissue Bank. A maximum Braak tau stage of 3 was used to filter out individuals with excessive Alzheimer’s pathology, thus ensuring that dementia in these individuals arose from α-synucleinopathy. PD without cognitive impairment was defined either by (i) a lack of evidence of positive cognitive features, such as memory impairment, executive dysfunction and visuo-spatial dysfunction in retrospective clinical case notes, or (ii) where positive cognitive features were reported present cognitive impairment was ruled out based on objective cognitive testing, or were proven to be as adverse effects of medication. Additionally, where possible, individuals were selected based on post-mortem interval less than 24 hours to ensure optimal tissue quality for nuclear extraction. In total, 7 PD, 7 PDD and 7 DLB individuals were selected, matched where possible for demographic and pathologic factors, along with 7 age-matched non-neurological control individuals. Control individuals were defined by a lack of clinical neurological features and no definitive pathological diagnoses. To ensure consistency, a cut-off of Braak tau stage 3 was also used for control individuals. The severity of α-synuclein pathology in the anterior cingulate was graded semi-quantitatively from 0-3 based on the validated scoring system from Alafuzoff *et al.* [34] For each individual, a tissue block of cortical grey matter from the anterior cingulate was sectioned at 80 µm thickness. Adjacent sections were subsequently used for bulk-tissue RNA isolation (2 sections per sample) or isolation of nuclei for single-nuclei RNA-sequencing. Clinical, pathological and sample measures for the cohort are available in **Supplementary Table 1**.

### Isolation of nuclei

Nuclei were isolated using buffers prepared as in Krishnaswami *et al.* [31], including nuclei isolation medium #1 (NIM1), nuclei isolation medium #2 (NIM2), Homogenisation Buffer (HB), 29% and 50% vol/vol iodixanol dilutions. Briefly, brain tissue sections were suspended in 800 µL HB and homogenised in a pre-cooled 2 mL dounce homogeniser, with five strokes of the loose pestle, followed by 10-15 strokes with the tight pestle. The homogenate was filtered through a BD Falcon tube with a cell strainer cap (35 µm) and centrifuged at 1000 g for 8 minutes. Thereafter, nuclei were subjected to an additional clean-up step (density gradient centrifugation), as detailed in Krishnaswami *et al.*, albeit with centrifugation of the layered nuclei/29% iodixanol solution at 13,000 *g* for 40 minutes at 4°C. The supernatant was carefully removed, and the nuclei pellet washed with PBS buffer (PBS + 1% BSA + 0.2 U/ml RNAseIn), filtered through a BD Falcon tube with a cell strainer cap, centrifuged at 500 *g* for 5 minutes at 4 °C and washed again. Nuclei were counted using a LUNA-FL Dual Fluorescence Cell Counter (Logos Biosystems, L20001) using Acridine orange dye to stain nuclei.

### Nuclei encapsulation and single-nucleus RNA-sequencing data generation

All samples were processed as per 10X Genomics Chromium Single Cell Reagent Kits Protocol, (chemistry: Single Cell 3’ v2). Following manufacturer’s guidelines, the samples were processed to target 10,000 nuclei per sample. Briefly, we performed 8 cycles of cDNA amplification and 14 cycles of final indexing PCR. cDNA concentrations were measured using Qubit dsDNA HS Assay Kit (ThermoFisher, Q32851), and cDNA and library preparations were assessed using the Bioanalyzer High-Sensitivity DNA Kit (Agilent, 5067-4627). All samples were pooled to equimolar concentration and sequenced together across twenty-eight lanes on an Illumina Hi-Seq 4000.

### Single-nucleus RNA-sequencing data processing

Sequenced reads were demultiplexed and processed using Cell Ranger (v 3.0.2) and thereafter mapped to the GRCh38 human reference genome using gene annotations from Ensembl v93 [81,82]. Across each of the 28 sequenced samples, reads mapped to primary transcripts were summarised as counts. Droplets containing nuclei were distinguished from empty droplets (containing ambient RNA) using the EmptyDrops algorithm, as implemented in the R package DropletUtils (v 1.6.1) [83]. An ambient profile threshold of 300 UMI was used to determine the background RNA content of the empty droplets. Thereafter we removed nuclei with > 5% mitochondrial content and genes expressed in < 5 nuclei. Once low-quality nuclei had been filtered out, the dataset was normalised using the `NormalizeData()` function in Seurat (v 3.2.0) [84]. The default normalising method used by Seurat (version 3) is a global-scaling normalisation method, “LogNormalize”. The method normalises the gene expression values in each cell (n) by multiplying n by the total expression of the cell (a size factor of 10,000 for each cell is used by default) and log-transforming the result. After this normalisation step, we used Seurat’s pipeline to cluster the nuclei. First, distances were calculated between two nuclei with similar gene expression patterns using Euclidean algorithm and edges were drawn. Second, a Louvain algorithm was used to cluster the nuclei. Finally, clustering was carried out using the `FindClusters()` function using 30 principal components (PCs) and a resolution parameter of 2. The clustered cells were tested to remove barcodes with more than 1 nuclei encapsulated in the droplet using DoubletFinder (v 2.0.2), with the expected proportion of doublets set at ∼7% [85].

### Cell-type identification

The remaining nuclei were visualised using a non-linear dimensionality reduction algorithm known as Uniform Manifold Approximation and Projection (UMAP, v 0.1.10) [86]. We then used the Wilcoxon rank sum test (FDR < 0.05) implemented in the Seurat function `FindAllMarkers()` to identify genes differentially expressed in one cluster compared with all other clusters. Cell types were assigned by testing genes differential to a particular cell-type for enrichment (Fisher’s exact test) for cell-type markers from two human single-cell datasets [32,87]. Nuclei classified as endothelial cells and pericytes were merged into one class referred to as vascular cells.

A joint graph of 205,498 nuclei from across all individuals from each of their respective filtered datasets (referred to as the panel of datasets) was generated using the R package, Clustering On Network Of Samples (Conos, v 1.1.2) [38]. This was done to bring panel datasets into a common expression space accounting for technical differences between datasets, which could be used for downstream cell-type-specific differential expression analyses between disease groups. buildGraph() was used to construct a graph with parameters for nearest neighbour parameters set at k=30, k.self=5, in space of 30 CPCA (common principal component). embedGraph() function was used to partition cells into 7 clusters for the 7-broad cell-types.

### Bulk-tissue RNA-sequencing data generation

RNA isolation was performed by the commercial company, BioXpedia A/S. Samples were lysed with QIAzol and RNA extracted using the RNeasy 96 Kit (Qiagen) with an optional on-membrane DNase treatment, as per manufacturer instructions. Samples were thereafter quantified by absorption on the QIAxpert (Qiagen) and their RNA integrity number (RIN) assessed using the Agilent 4200 Tapestation (Agilent). RIN ranged from 1.6-7.8, with a median of 6.5. Only samples derived from tissue-sections with a RIN ≥ 4.2 were included in downstream RNA sequencing. As a result, only 24 samples were sequenced (5 controls, 7 PD, 6 PDD and 6 DLB; **Supplementary Table 1**). 250 ng of total RNA was used as input for cDNA library construction with the TruSeq Stranded mRNA Sample Preparation Kit (Illumina), as per manufacturer instructions. To minimise read mis-assignment in downstream sample de-multiplexing, xGen UDI-UMI Adapters (Integrated DNA Technologies, Inc.) were used. Libraries were multiplexed on the NovaSeq S2 Flow Cell (the same 24 libraries were run across both lanes) for paired-end 100 bp sequencing on the NovaSeq 6000 Sequencing System (Illumina) to obtain an average read depth of ∼180M paired-end reads per sample.

### Bulk-tissue RNA-sequencing data processing

Fastp (v 0.20.0), a fast all-in-one FASTQ pre-processor, was used for adapter trimming, read filtering and base correction [88]. Fastp default settings were used for quality filtering and base correction. Processed reads were mapped to the GRCh38 human reference genome via STAR (v 2.7.0a) using gene annotations from Ensembl v97 [81,82]. Multi-sample 2-pass mapping was used, wherein two rounds of mapping were performed to improve the sensitivity of novel splice junction detection. ENCODE standard options for long RNA-seq were used, with the exception of (i) `--outFilterMultimapNmax`, which was set to 1, thus retaining only uniquely mapped reads, and (ii) `--alignSJDBoverhangMin`, which was set to the STAR default of a minimum 3 bp overhang required for an annotated spliced alignment. Processed reads were also quantified with Salmon (v 0.14.1) using the mapping-based mode, with sequence-specific, fragment GC-content and positional bias correction options enabled (`--seqBias`, `--gcBias`, `--posBias`) [89]. A *decoy-aware* transcriptome file based on GRCh38 and Ensembl v97 was generated using MashMap2 (v 2.0) [90] and used as a reference together with the appropriate option for the sequencing library type (`--libType ISF`).The R package *tximport* (v 1.14.2) was used to transform Salmon transcript-level abundance estimates to gene-level abundance estimates [91]. Genes found to overlap ENCODE blacklist regions were removed from downstream analyses (**Key resources**) [92]. Pre-alignment quality control metrics were generated using Fastp and FastQC (v 0.11.8) [93], and post-alignment quality control metrics using RSeQC (v 2.6.4) [94]. Pipeline source code can be found in https://github.com/RHReynolds/RNAseqProcessing.

### Processing of PD case-control replication dataset

Replication of several downstream bulk-tissue RNA-sequencing analyses were performed using a PD case-control bulk-tissue RNA-sequencing dataset provided by Dumitriu *et al.* [44] and processed for re-use by recount2 [95]. The dataset was accessed via recount2 (recount accession ID: SRP058181). The original study contained RNA-sequencing of prefrontal cortical samples (Brodmann Area 9) derived from 44 control individuals and 29 individuals with PD. Paired-end 101-bp sequencing was applied to each sample, with a mean depth of 83.3 million read pairs per sample. All samples were of a reasonably high quality with RIN values ranging from 5.8-9.1 and a median of 7.6. Accessed samples were checked for any mismatch between the reported sex of brain donors and the sex as determined by the expression of sex-specific genes (*XIST* and *DDX3Y*). As a result, one control sample was removed (recount sample ID: SRR2015746; study sample ID: C0061); the sample was reported to be male, but notable expression of *XIST* was observed. Further, as sample demographics from the original study included whether PD patients were diagnosed with dementia, the 29 PD cases were split into those with and without dementia (PD, n = 18; PDD, n = 11).

### Deconvolution

Cell-type proportions in bulk-tissue RNA-sequencing samples were estimated using Scaden (v 0.9.2), a deep-learning-based deconvolution algorithm [43]. Unlike linear-regression-based deconvolution algorithms, Scaden does not require cell-type-specific expression profiles. Instead, Scaden trains on artificial bulk-tissue RNA-sequencing samples simulated from tissue-specific single-cell RNA-sequencing data, after which the model is used to predict cell-type proportions from real bulk-tissue RNA-sequencing samples. In this study, training data was generated separately for each individual with paired single-nucleus RNA-and bulk-tissue RNA-sequencing, allowing Scaden to capture cross-subject heterogeneity. This yielded a total of 24,000 artificial bulk-tissue RNA-sequencing samples (1,000 samples per subject). Prior to generation of training data, single-nucleus RNA-sequencing counts per cell were normalised using the total counts over all genes, ensuring that every cell had the same total count after normalisation. Thereafter, artificial bulk-tissue RNA-sequencing samples were simulated using the Scaden `bulk_simulation.py` script, which sub-samples cells from input single-nucleus RNA-sequencing data and then aggregates expression across sub-sampled cells. Here, 1,000 cells were used per simulated sample. Artificial bulk-tissue RNA-sequencing samples were combined and stored in a h5ad file, using the Scaden `create_h5ad_file.py` script. To ensure generated training data and bulk-tissue RNA-sequencing samples (in the form of counts normalised by library size) for prediction shared the same features (genes) and feature scale, both datasets were pre-processed with `scaden process` (the two datasets shared a total of 13,191 genes following processing). Following this, each of the three Scaden ensemble models was independently trained (`scaden train`) for 5,000 steps, as recommended by the developers to prevent overfitting, using the default values for batch size and learning rate[43]. Finally, predictions for cell-type proportions were made with `scaden predict`.

Replication of predicted cell-type proportions was performed using a second independent PD case-control dataset accessed from recount2 (see **Processing of PD case-control replication dataset**). As the Scaden algorithm requires that training data and prediction data have a perfect overlap of features, it was necessary to re-perform pre-processing with `scaden process` (using library-normalised counts from the replication dataset; the two datasets shared a total of 14,094 genes following processing) and to train a new model (using the same parameters as previously). In both datasets, significant differences in cell-type proportions between disease groups were a two-sided Wilcoxon rank sum test, with FDR-correction for multiple testing.

### Bulk-tissue RNA-sequencing covariate selection

Sources of variation in bulk-tissue RNA-sequencing data were identified using principal component analysis (PCA) performed on gene-level expression filtered to include only genes with count > 0 in all samples (28,692 genes) and transformed with DESeq2’s `vst()`, which applies a variance stabilising transformation. RIN and age of death were significantly correlated with the first and second PC, respectively. Furthermore, cell-type proportions for excitatory and inhibitory neurons, microglia and astrocytes were significantly correlated with the first, third and fourth PC, respectively. Thus, the final model for differential expression and splicing (referred to as the “cell-type- and covariate-corrected” model) consisted of the disease group and the top 4 PCs (which collectively explained 52.6% of the total variance).

To explore the effect of accounting for cell-type proportions, vst-transformed gene expression was batch-corrected using the final “cell-type- and covariate-corrected” model or a minimised “covariate-corrected” model consisting of disease group, age of death, RIN and sex. Samples were thereafter plotted by their first two principal components to determine how well disease groups separated (**Supplementary Figure 6**). Batch correction was performed using the `removeBatchEffect()` function from the R package, limma (v 3.42.2) [96]. Prior to correction, covariates to be used in the model were scaled to ensure that variables that are measured on different scales (e.g. age of death vs RIN) are comparable.

As in the original study [44], the final model for the replication dataset (see **Processing of PD case-control replication dataset**) included disease group and the covariates age of death, RIN and post-mortem interval (PMI). In addition, cell-type proportions for all cell types were included in the final model, as these were significantly correlated with several of the top 8 PCs.

### Differential gene expression

#### Single-nucleus RNA-sequencing

We used Model-based Analysis of Single-cell Transcriptomics (MAST, v 1.12.0), a method specifically designed to carry out differential expression analysis on our single-nucleus RNA-sequencing data [97]. MAST is a two-part, generalized linear model. The first part of the model uses logistic regression to model whether a gene is expressed i.e. the discrete rate of expression of each gene over the background of other transcripts. The second part of the model models the level of expression (conditional on whether a gene is expressed in a cell) using a Gaussian linear model. Information from both parts of the model are combined to model changes in gene expression levels and with control for multiple sources of variation such as cell-cell variation. MAST also models the cellular detection rate, which is defined as the fraction of genes that are detectably expressed in each cell. The cellular detection rate acts as a substitution for both technical and biological factors such as dropout, cell volume and other extrinsic factors that could influence gene expression. Controlling for the cellular detection rate improves the sensitivity (true positive rate) and specificity (true negative rate) of MAST in the presence of confounding between the cellular detection rate and true biological signals.

To perform differential expression, cell-type-specific nuclei from each of the 28 filtered sample count matrices (see **Single-nucleus RNA-sequencing data processing**) were merged to create 7 cell-type count matrices. Genes that were expressed in ≤ 3 nuclei were removed from the analysis. Following this, differential expression analysis was performed separately for each cell type, across all pairwise combinations of the disease groups (n = 6). A likelihood ratio test was used, with age of death, post-mortem interval (PMI), and sex included as covariates. Genes with FDR < 0.05 and absolute fold-change > 1.5 were considered significant.

#### Bulk-tissue RNA-sequencing

Bulk-tissue differential gene expression was assessed using the DESeq2 R package (v 1.26.0) and gene-level expression filtered to include only genes with count > 0 in all samples (28,692 genes) [98]. With one exception (the maximum number of iterations allowed for convergence, `maxit = 1000`), default parameters were used, including the default Wald test of significance. Differentially expressed genes were identified in a pairwise manner, controlling for covariates identified using gene-level expression (see **Bulk-tissue RNA-sequencing covariate selection**). Multiple testing was performed by FDR-correction, with a cut-off of FDR < 0.05 applied for significance.

### Differential splicing analysis

Differential splicing was assessed using Leafcutter (v 0.2.8), which detects splicing variation using sequencing reads with a gapped alignment to the genome (here, termed junction reads) [60]. Junction reads, which are presumed to represent intron excision events, are used to quantify intron usage across samples without any reliance on existing reference annotation. Importantly, Leafcutter does not estimate isoform abundance or exon inclusion levels, but rather captures changes in local splicing events through construction of intron clusters, wherein overlapping introns are connected by the splice junction(s) they share. As input, splice junctions outputted by STAR (`SJ.out.tab`) were first filtered to remove any regions that overlap ENCODE blacklist regions (**Key resources**) [92] and thereafter converted to the `.junc` files used by Leafcutter for intron clustering. The conversion was performed using custom R code (`convert_STAR_SJ_to_junc()` in https://github.com/RHReynolds/RNAseqProcessing). Intron clusters were defined using Leafcutter’s `leafcutter_cluster.py` with thresholds ensuring that (i) introns supported by < 30 junction reads across all 24 samples or < 0.1% of the total number of junction read counts for the entire cluster and (ii) introns of more than 1Mb were removed. This yielded a total of 43,544 clusters encompassing 152,298 introns that were used for further analysis. Differentially spliced (DS) clusters were identified in a pairwise manner, controlling for covariates identified using gene-level expression (see **Bulk-tissue RNA-sequencing covariate selection**), and annotated to genes using exon files generated from GRch38 Ensembl v97 (with the Leafcutter helper script `gtf_to_exons.R`). As per Leafcutter default filters, only introns detected in ≥ 5 samples were tested and an intron cluster was only tested if detected in ≥ 3 individuals in each comparison group with an overall coverage of ≥ 20 junction reads. P-values were FDR-corrected for multiple testing and an intron cluster and its overlapping gene were considered differentially spliced if (i) FDR < 0.05 and (ii) the intron cluster contained at least one intron with an absolute delta percent-spliced-in value (|ΔPSI|) ≥ 0.1. The latter filter was applied to improve the specificity of Leafcutter [99].

### Annotation of differential splicing events

Introns within intron clusters were annotated using `annotate_junc_ref()`from the R package **D**etecting **A**berrant **Sp**licing **E**vents from **R**NA-sequencing (dasper, v 1.1.4) [100], which categorises junctions based on (i) whether the junction is present within the entire set of annotated introns or (ii) whether both, one of, or neither the donor and acceptor splice site precisely overlap the boundary of a known exon. For both checks, Ensembl v97 was used. When defining and clustering introns, `leafcutter_cluster.py` adds 1 bp to the end of a junction read; thus, to ensure optimal mapping to reference annotation, 1 bp was removed from all intron ends prior to use of `annotate_junc_ref()` using custom code (`convert_leafcutter.R` from https://github.com/RHReynolds/LBD-seq-bulk-analyses). Junctions (and the introns they represent) were then classified into one of the following categories: annotated, novel exon skip, novel combination, novel acceptor, novel donor, ambiguous gene and unannotated (“none”) (**Supplementary Figure 13**). Annotated junctions are those that match the boundaries of an existing intron. Unannotated junctions have neither end overlapping a known exon. Novel acceptors and novel donors are junctions where one end (acceptor or donor) matches the boundary of a known exon. Novel exon skip and novel combination junctions have both ends overlapping known exon boundaries, which are not part of the set of annotated introns. They are distinguished by whether their start or end overlaps exons derived from the same transcript. That is, for an event to be a novel exon skip, both the start and end must overlap an exon contained in the same transcript, whereas to be a novel combination, the start and end overlap exons are from different transcripts. Junctions that mapped to more than one gene (“ambiguous gene”) were not considered in downstream analyses.

### Gene set enrichment

#### Functional enrichment of cell-type-specific differentially expressed genes

Functional term enrichment analysis for cell-type-specific differentially expressed genes from each pairwise comparison was performed using the overrepresentation analysis module from the R package implementation of WEB-based Gene SeT AnaLysis Toolkit (WebGestaltR, v 0.4.4) [101]. Two separate analyses were performed using (i) only non-redundant Gene Ontology (GO) terms (which are generated by selecting the most general terms in each branch of the GO directed acyclic graph structure from all terms with 20-500 genes) and (ii) 46 biological pathways associated with PD risk in a large-scale pathway-specific polygenic risk analysis [46]. For both analyses, default values for WebGestalt parameters were used, which include a minimum and maximum overlap of 10 and 500, respectively. FDR-correction for multiple testing was performed, and significant pathways were those with FDR < 0.05.

#### Functional enrichment of differentially spliced genes

Gene set enrichment for GO terms was performed using `enrichGO()` and `clusterCompare()` from clusterProfiler (v 3.14.3), which permit GO enrichment analysis (based on a hypergeometric distribution) and comparison across multiple gene lists [102]. Two separate analyses were run using (i) all differentially spliced genes (FDR < 0.05, |ΔPSI| >= 0.1) across each pairwise comparison in the discovery dataset and (ii) genes overlapping validated intron clusters with ≥ 1 intron that shared the same direction of effect. In both analyses, default parameters were used; these included FDR-correction for multiple testing and filtering for terms with FDR < 0.05.

Functional enrichment of genes associated with bulk-tissue gene expression principal components Genes contributing to PC1, following batch correction of cell-type proportions (as described in **Bulk-tissue RNA-sequencing covariate selection**), were extracted using `get_pva_var()` from the R package, factoextra (v 1.0.7). The top 100 genes contributing to gene-expression-derived PC1 were used for gene set enrichment with `enrichGO()` from clusterProfiler [102]. Default parameters were used, which included FDR-correction for multiple testing and filtering for terms with FDR < 0.05.

#### Visualisation of GO term overlaps between analyses

Overlapping GO-derived pathway enrichments from each of the three analyses (i.e. single-nucleus differential expression, bulk-tissue differential splicing, and gene expression contributions to bulk-tissue PC1) were visualised using the ComplexHeatmap R package (v 2.7.7) [103]. Pathways from all three analyses were filtered to include only those that were shared across more than one type of analysis. Pathways were encoded by a binary 1 and 0 for present and absent, respectively, permitting clustering of gene sets by Pearson correlation. Gene sets derived from differential splicing were collapsed across our own dataset and the validation dataset, resulting in one gene set per pairwise comparison. Likewise, gene sets derived from up- and down-regulated single-nucleus DE gene sets were collapsed across cell types, resulting in 7 gene sets per pairwise comparison.

#### Reduction of GO terms using semantic similarity

To reduce redundancy across GO-derived pathway enrichment analyses derived from various analyses (i.e. single-nucleus differential expression, bulk-tissue differential splicing, genes contributing to bulk-tissue PC1), two steps were taken. First, GO terms were filtered to exclude terms with ≥ 20 genes or ≤ 2000 genes. Second, semantic similarity of all enriched GO terms was calculated using `mgoSim()` from the GOSemSim R package (v 2.17.1) [104] and a graph-based measure of semantic similarity (`measure = “Wang”`) [105]. Thereafter, `reduceSimMatrix()` from the rrvgo R package (v 1.1.4) was used to reduce terms [106]. This function reduces terms by generating a distance matrix from the semantic similarity scores, which is hierarchically clustered using complete linkage (a “bottom-up” clustering approach). Both steps were combined into the function `go_reduce`(), available at: https://github.com/RHReynolds/rutils. The hierarchical tree was then cut at a threshold of 0.9 (leading to fewer groups), and the term with the highest semantic similarity score was used to represent each group of terms. This reduction was performed separately for each of the three analyses.

#### Cell-type enrichment of differentially spliced genes

Expression-weighted cell-type enrichment (v 0.99.2) was used to determine whether differentially spliced genes demonstrate higher expression in certain cell types than would be expected by chance [107]. EWCE requires two inputs: a gene list and gene cell-type specificity values derived from single-cell/nucleus data (here, termed a specificity matrix). Two sets of gene lists were run. The first set of gene lists included the top 100 differentially spliced genes (FDR < 0.05, |ΔPSI| >= 0.1, ranked by p-value) across each pairwise comparison in the discovery dataset. In the case where a gene had multiple significant intron clusters, the most significant cluster with the highest |ΔPSI| was used for ranking. The second set of gene lists included genes overlapping validated intron clusters with ≥ 1 intron that shared the same direction of effect. Both sets of gene lists were run together with gene cell-type specificity values separately derived from each disease group (i.e. Control, PD, PDD and DLB); specificity matrices were generated for cell types in each disease group using the `generate.cell.data()` function of the EWCE package. For each combination of gene list and specificity matrix, 100,000 bootstrap replicates were used. Transcript length and GC-content biases were controlled by selecting bootstrap replicates with comparable properties to the target gene lists. Data are displayed as standard deviations from the mean, which indicate the distance of the mean expression of the target gene list from the mean expression of the bootstrap replicates.

### RNA-binding protein binding motif analysis

#### Generating sequences

Two sets of sequences were generated per pairwise comparison. These sets included all differentially spliced introns (FDR < 0.05, |ΔPSI|) and non-differentially spliced introns (FDR > 0.05), as defined by their 5’ and 3’ proximal intronic regions (500 nucleotides of proximal intron and 50 nucleotides of exon flanking the 5’ and 3’ splice sites). A 5’ or 3’ splice site could be associated with more than one intron (e.g. in the case of two introns with the same 5’ splice site, but varying 3’ splice sites), and thus could be associated with more than one |ΔPSI| value. In these cases, the highest |ΔPSI| was assigned to the proximal intronic region.

#### Enrichment of RBP binding motifs

The position weight matrices (PWMs) of RBP binding motifs in humans were collected from the ATtRACT database (v 0.99β) [61]. Motifs < 7 nucleotides in length and with a quality score of < 1 were removed to reduce false positives in the motif matches (quality score estimates the binding affinity between RBPs and binding sites). Furthermore, to remove redundancy between multiple motifs for one RBP, the longest available motif was selected. Finally, RBPs that had a median TPM of 0 in GTEx (v 8) anterior cingulate cortex samples were removed (e.g. RBMY1A1) [108]. This resulted in 82 unique PWMs, which were used to identify enrichment of RBP binding motifs. Analysis of Motif Enrichment (AME, v 5.1.1) [109] was used with default parameters (`--scoring avg`) to compare enrichment of RBP binding motifs between differentially spliced and non-differentially spliced proximal intronic regions. RBP binding motifs with an enrichment-optimised and Bonferroni-adjusted p < 0.05 were considered to be significantly over-represented in differentially spliced proximal intronic regions compared with non-differentially proximal intronic regions.

### Integration with GWAS

To test for enrichment of genetic association of a gene set to a trait we employed two orthogonal methods, Hi-C-coupled Multi-marker Analysis of GenoMic Annotation (H-MAGMA) [51] and stratified LD score regression (sLDSC) [52]. Both methods were run with two sets of annotations: (i) the top 10% most cell-type-specific genes, as determined using specificity values derived from EWCE (see **Cell-type enrichment of differentially spliced genes**) and (ii) cell-type-specific differentially expressed genes (FDR < 0.05, |log_2_(fold change)| > log_2_(1.5)). These annotations were run with 3 genome-wide association studies (GWASs), including Alzheimer’s disease (AD), Parkinson’s disease (PD) and Parkinson’s disease Age of Onset (PD AOO) (**Table 1**) [27,56,57] In both analyses, p-values were FDR-corrected for the number of cell types tested.

**Table 1.** Summary of GWAS datasets. AD, Alzheimer’s disease; PD, Parkinson’s disease.

| Disease | First author, Year | N cases | N controls | PMID | Reference |
| --- | --- | --- | --- | --- | --- |
| AD | Jansen, 2019 | 71,880 | 383,378 | 30617256 | [57] |
| PD –<br>risk | Nalls, 2019 (excluding<br>23andMe contributions) | 33,674<br>(18,618 proxy cases<br>from UK Biobank) | 449,056 | 31701892 | [27] |
| PD –<br>age of<br>onset | Blauwendraat, 2019 | 17,415 |  | 30957308 | [56] |

### H-MAGMA

Hi-C-coupled MAGMA (H-MAGMA) (v 1.08b of MAGMA[110]) was used to carry out gene-set enrichment analysis using three GWAS summary statistics. Gencode v26 (**Key resources**) was used to assign exonic SNPs and promoter SNPs, which is defined as 2kb upstream of the transcription start site (TSS), to their target genes based on their genomic location. Chromatin interactions to exons and promoters generated from Hi-C performed on adult dorsolateral prefrontal cortex, were used to assign intergenic and intronic SNPs to their cognate genes [51]. Gene-level association statistics were computed using window coordinates of 10kb downstream and 35kb upstream.

### sLDSC

Stratified LDSC (v 1.0.1) was used to test whether cell-type specific DE genes or the top 10% most cell-type-specific genes contributed to the common SNP heritability of AD, PD or PD AOO [111,112]. To ensure gene lists were sufficiently large, only gene lists with more than 20 genes were run. Gene coordinates (Ensembl v97, GRCh38) were extended by 100kb upstream and downstream of their transcription start and end site, in order to capture regulatory elements that might contribute to disease heritability [112]. All annotations were constructed in a binary format (1 if the SNP was present within the annotation and 0 if not), using all SNPs with a minor allele frequency > 5%. Annotations were then added individually to the baseline model of 53 annotations provided by Finucane et al. (v 1.2, GRCh38), comprising genome-wide annotations reflecting genetic architecture. As annotations and the baseline model were mapped to GRCh38, all GWAS summary statistics were converted from GRCh37 to GRCh38 using the R implementation of the LiftOver tool, which is available from the rtracklayer package (v 1.46.0) [113]. HapMap Project Phase 3 (HapMap3) SNPs and 1000 Genomes Project Phase 3 European population SNPs were used for the regression and LD reference panels, respectively [114,115]. The MHC region (chr6: 25000000 – 34000000, GRCh38) was excluded from all analyses owing to the complex and long-range LD patterns in this region. For all stratified LDSC analyses, we report a one-tailed p-value (coefficient p-value) based on the coefficient z-score outputted by stratified LDSC. A one-tailed test was used as we were only interested in annotation categories with a significantly positive contribution to trait heritability, conditional upon the baseline model.

### Key resources

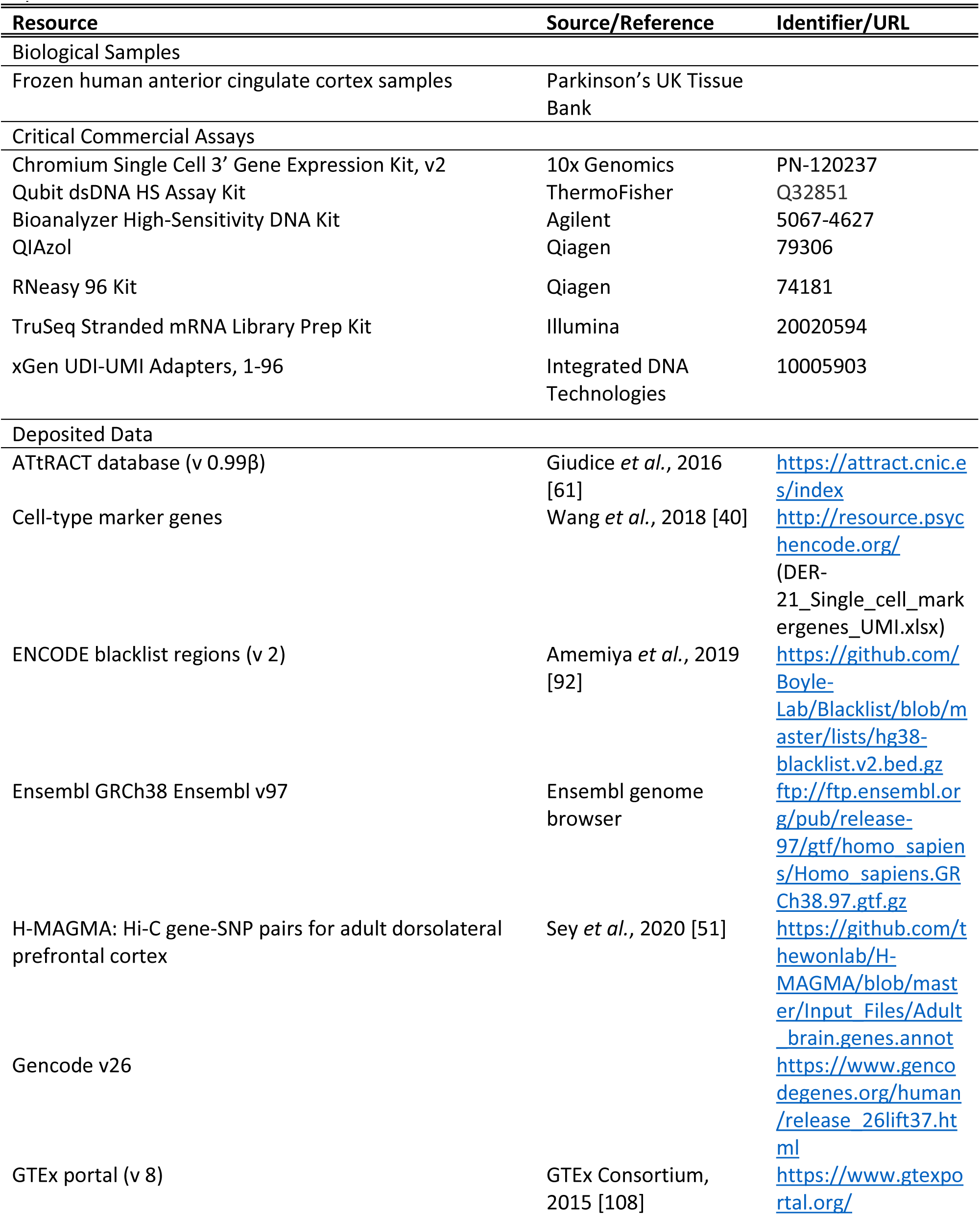
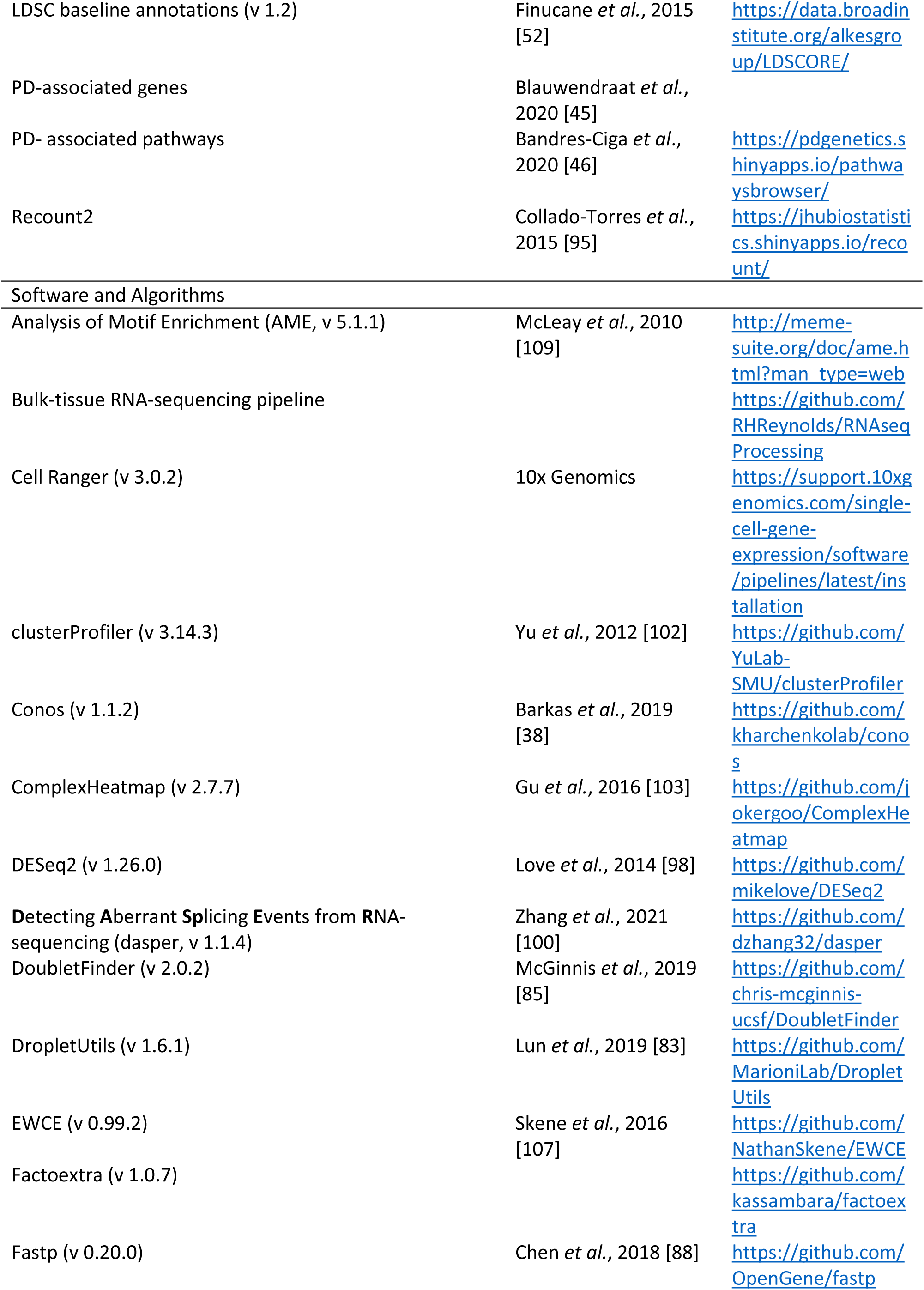
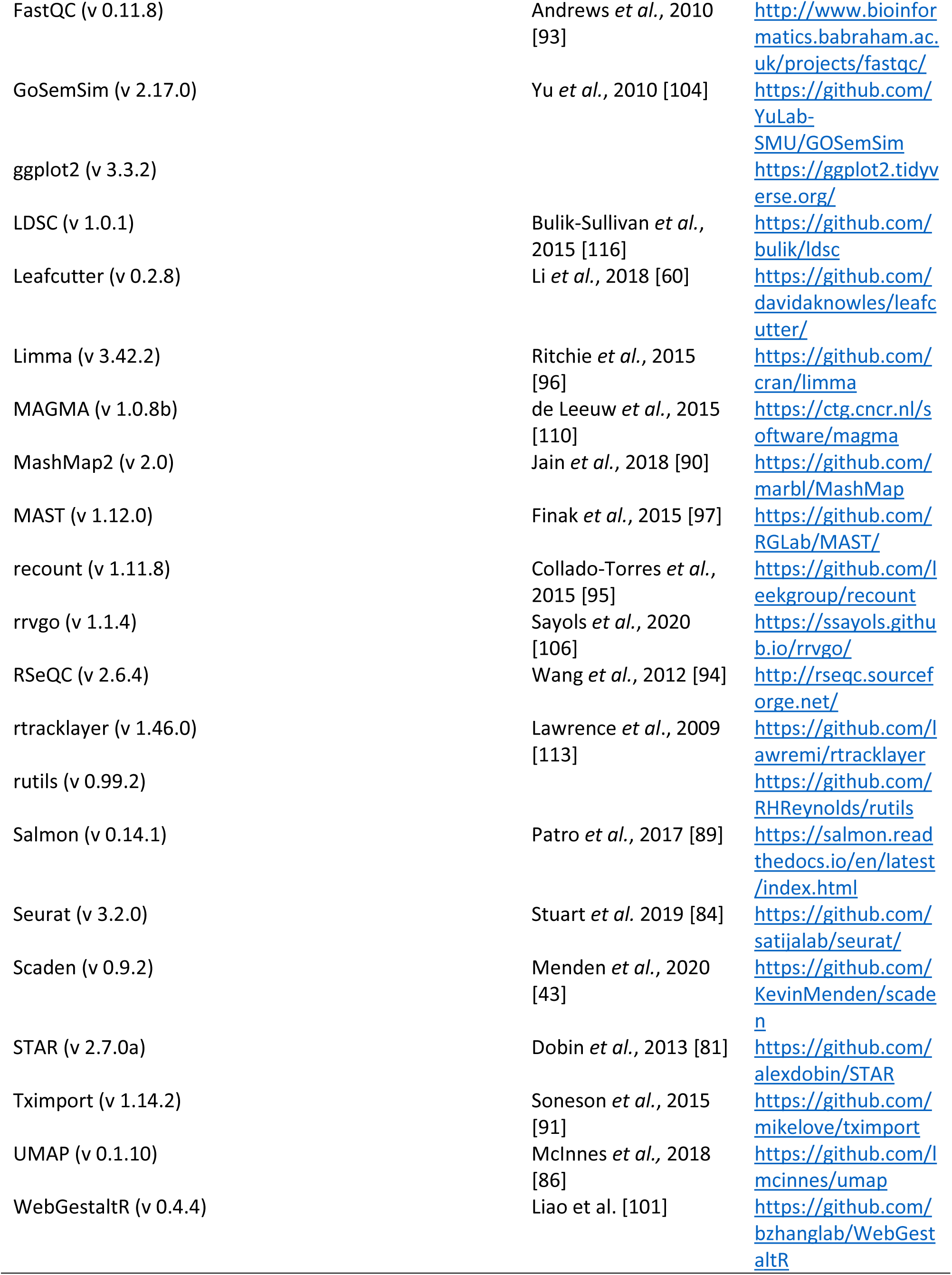

## Declarations

### Ethics approval

Ethics approval for the work carried out on the tissue from the Multiple Sclerosis and Parkinson’s Tissue Bank was given by Wales REC3 ethic committee, REC reference 18/WA/0238.

### Code availability

Code used to process and analyse bulk-tissue RNA-sequencing data, to generate sLDSC outputs, and to generate figures for the manuscript will be available upon publication at: https://github.com/RHReynolds/LBD-seq-bulk-analyses. Code used to process and analyse single-nucleus RNA-sequencing data and to generate H-MAGMA outputs will be available upon publication at: https://github.com/johnsonlab-ic/LBD-seq-singlenuc-analyses. All other open source software used in this paper is available for all tools used (see **Key resources**).

### Data availability

Bulk-tissue RNA-sequencing data will be made available on EGA upon publication. Single-nucleus RNA-sequencing data will be made available on GEO upon publication.

## Acknowledgements

RF, DO, MRJ and PS were supported through the UKRI Medical Research Council (MRC grant code, DO: MR/N008219/1; RF, MRJ and PS: MR/S02638X/1). MRJ was also separately supported through the Imperial College NIHR Biomedical Research Centre (BRC) Scheme. RHR was supported through the award of a Leonard Wolfson Doctoral Training Fellowship in Neurodegeneration and through the Signe og Peter Gregersens Mindefond. AS was supported through the UK Dementia Research Institute. SAGT acknowledges support from a Junior 1 award from the Fonds de recherche du Québec - Santé (FRQS). JH was supported through the UKRI Medical Research Council (MRC grant code: MR/N026004/), the UK Dementia Research Institute, The Wellcome Trust (202903/Z/16/Z), the Dolby Family Fund, and the NIHR. PMM was supported through the Imperial College NIHR Biomedical Research Centre (BRC) and the UK Dementia Research Institute and gratefully acknowledges personal funding from the Edmond Safra Foundation and Lily Safra. He is an NIHR Senior Investigator. SG is director of the Parkinson’s UK Tissue Bank, funded by Parkinson’s UK, a charity registered in England and Wales (258197) and in Scotland (SC037554). MR was supported through the award of a UKRI Medical Research Council Clinician Scientist Fellowship (MRC grant code: MR/N008324/1).

## Competing interests

RF, RHR, AS, BT, SAGT, JH, SG, DO, MRJ, PS and MR declare that they have no relevant financial or non-financial interests to disclose. PMM has received honoraria or consulting fees from Biogen, Novartis, Ipsen Pharmaceuticals, NodThera and Celgene. He receives research funding from Biogen, Merck, Celgene and Bristol Myers Squibb.

## Author contributions

RF processed and analysed single-nucleus RNA-sequencing data and assisted with preparation of the manuscript. RHR processed and analysed bulk-tissue RNA-sequencing data, integrated bulk-tissue and single-nucleus RNA-sequencing data, and together with MR, prepared the first draft of the manuscript. AS generated tissue sections, isolated nuclei and was involved in quality control of single-nucleus and bulk-tissue RNA-sequencing data generation. BT assisted with access to tissue samples and performed the neuropathological assessment of all samples. SAGT assisted with genetic enrichment analyses. SG enabled access to tissue samples and supervised their neuropathological assessment. MRJ and PS supervised processing and analysis of single-nucleus RNA-sequencing data and genetic enrichment analyses. MR supervised processing and analysis of bulk-tissue RNA-sequencing data and integration with single-nucleus RNA-sequencing data. AS, JH, PMM, SG, DO, MRJ, PS and MR conceived and designed the study. All authors contributed to the critical review of the manuscript.

## Supplementary Figures

**Supplementary Figure 1.**
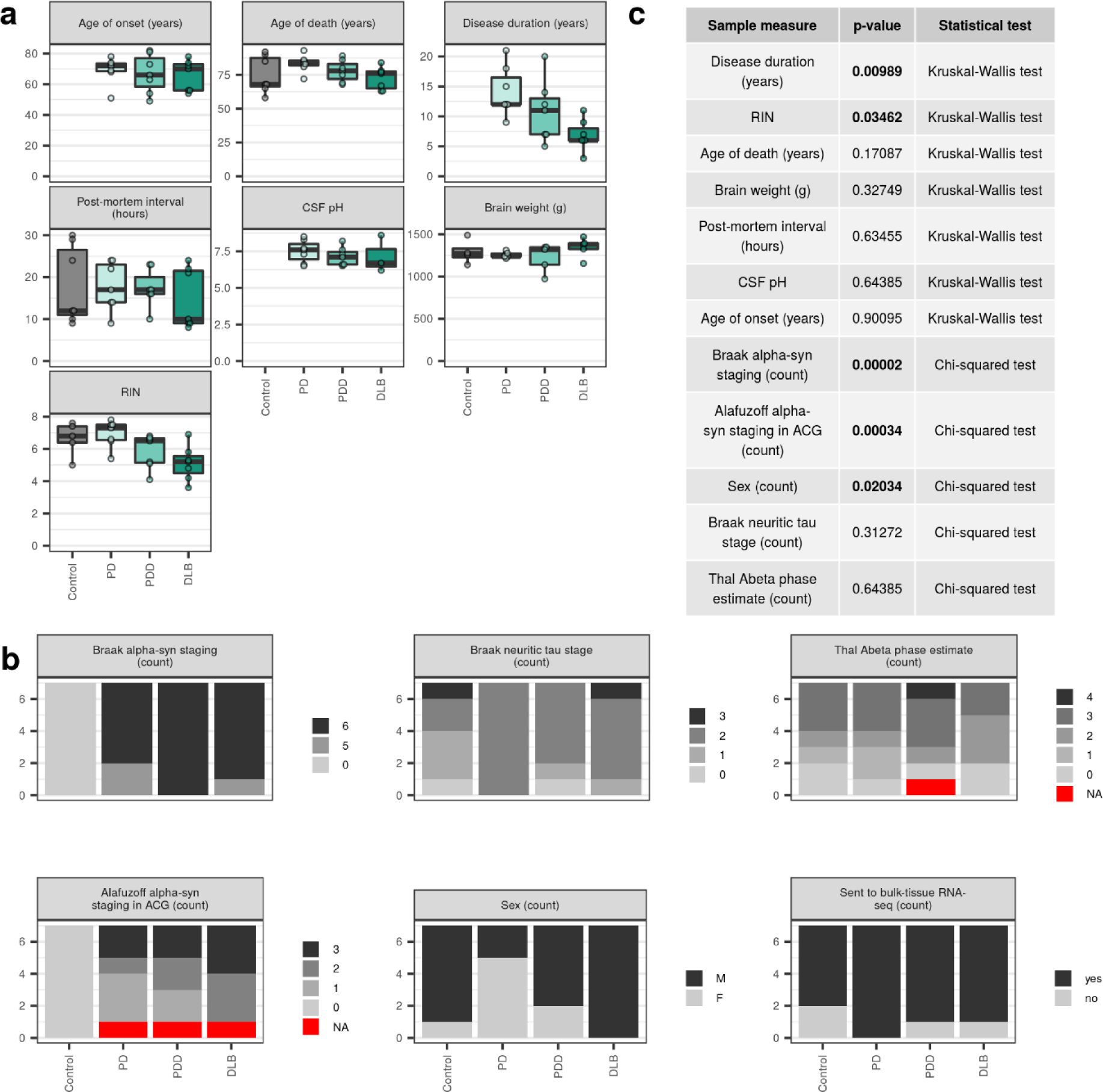
Subject demographics, sample information and pathological measures. Continuous **(a)** and categorical **(b)** subject demographics, sample measures and pathological measures are shown for each disease group. **(c)** Significant differences in groups were tested using either the Kruskal-Wallis rank sum test (for continuous variables) or the Chi-squared test (for categorical variables). Significant differences (p < 0.05) are shown in a bold face. All measures per individual are available in **Supplementary Table 1**.

**Supplementary Figure 2.**
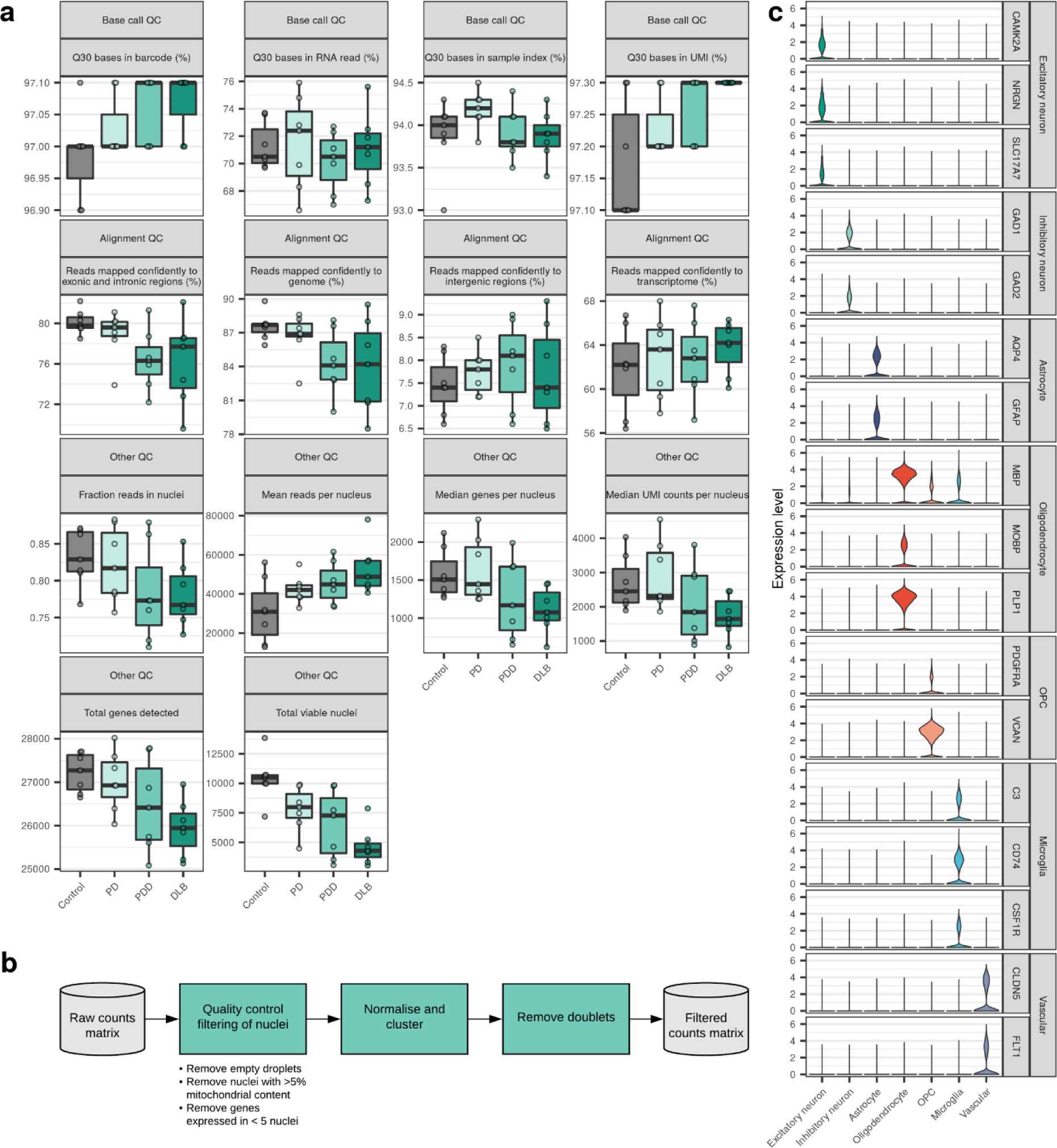
Single-nucleus RNA-sequencing metrics, quality control and cell-type classification. **(a)** Single-nucleus sequencing metrics from Cell Ranger, grouped by the type of quality control (QC) metric they represent i.e. base call, alignment or other. All metrics per sample are available in **Supplementary Table 1**. **(b)** Workflow illustrating the steps taken to filter the raw data to only include true nuclei of high quality. These steps were independently applied across each of the 28 sequenced samples. **(c)** Violin plots of expression values (y-axis) for known cell-type marker genes in each of the cell-type clusters identified (x-axis).

**Supplementary Figure 3.**
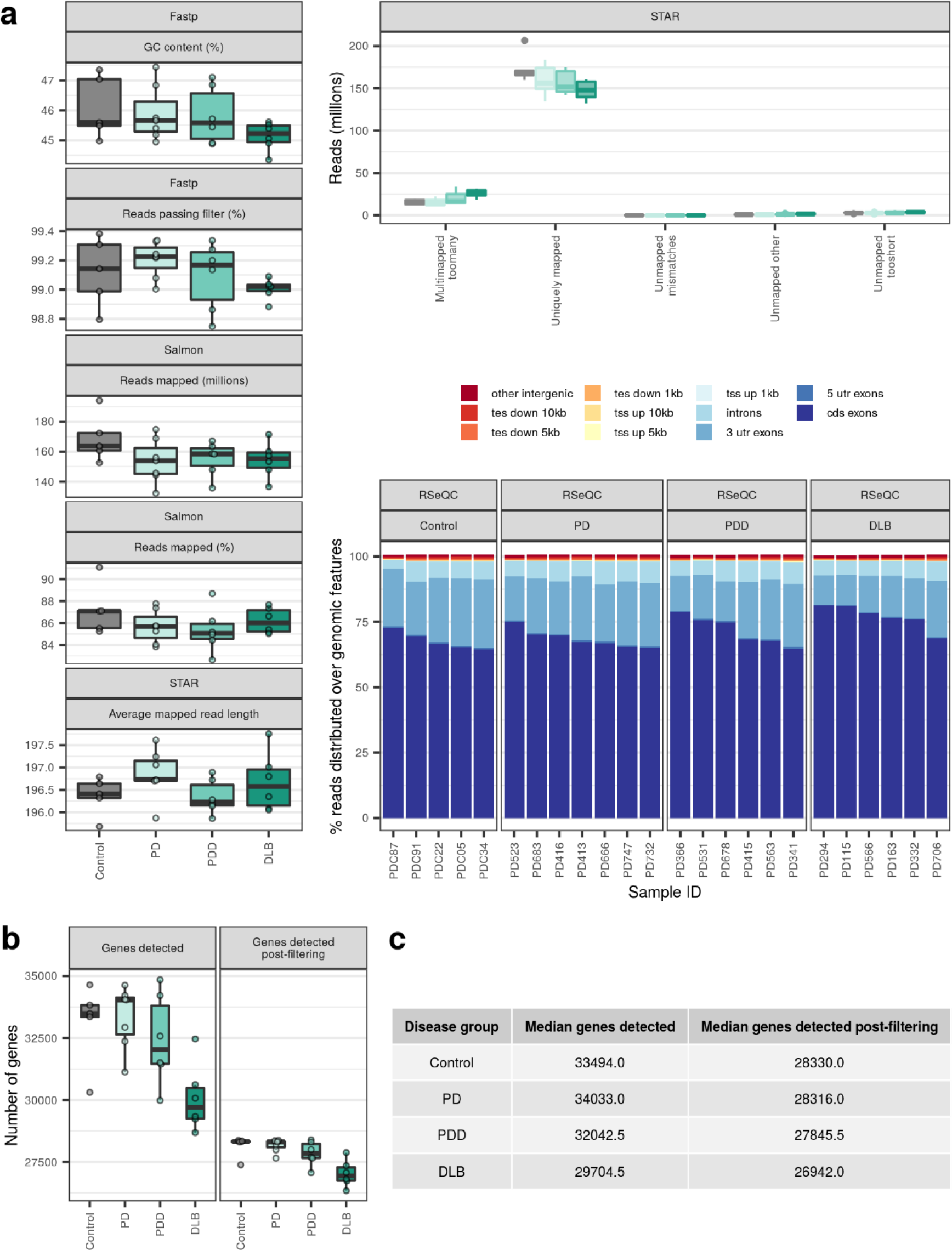
Bulk-tissue RNA-sequencing metrics. **(a)** Bulk-tissue RNA-sequencing metrics from Fastp, STAR, Salmon and RSeQC. **(b)** Number of genes detected in bulk-tissue RNA-sequencing of each sample before and after filtering. Detection of a gene before filtering was defined as a count > 0 in at least one sample across a disease group, while detection of a gene after filtering was defined as a count > 0 in all samples across a disease group. Only genes that were detected after filtering were used for bulk RNA-sequencing differential gene expression analyses. **(c)** Descriptive statistics for **(b)**. All metrics per sample are available in **Supplementary Table 1**.

**Supplementary Figure 4.**
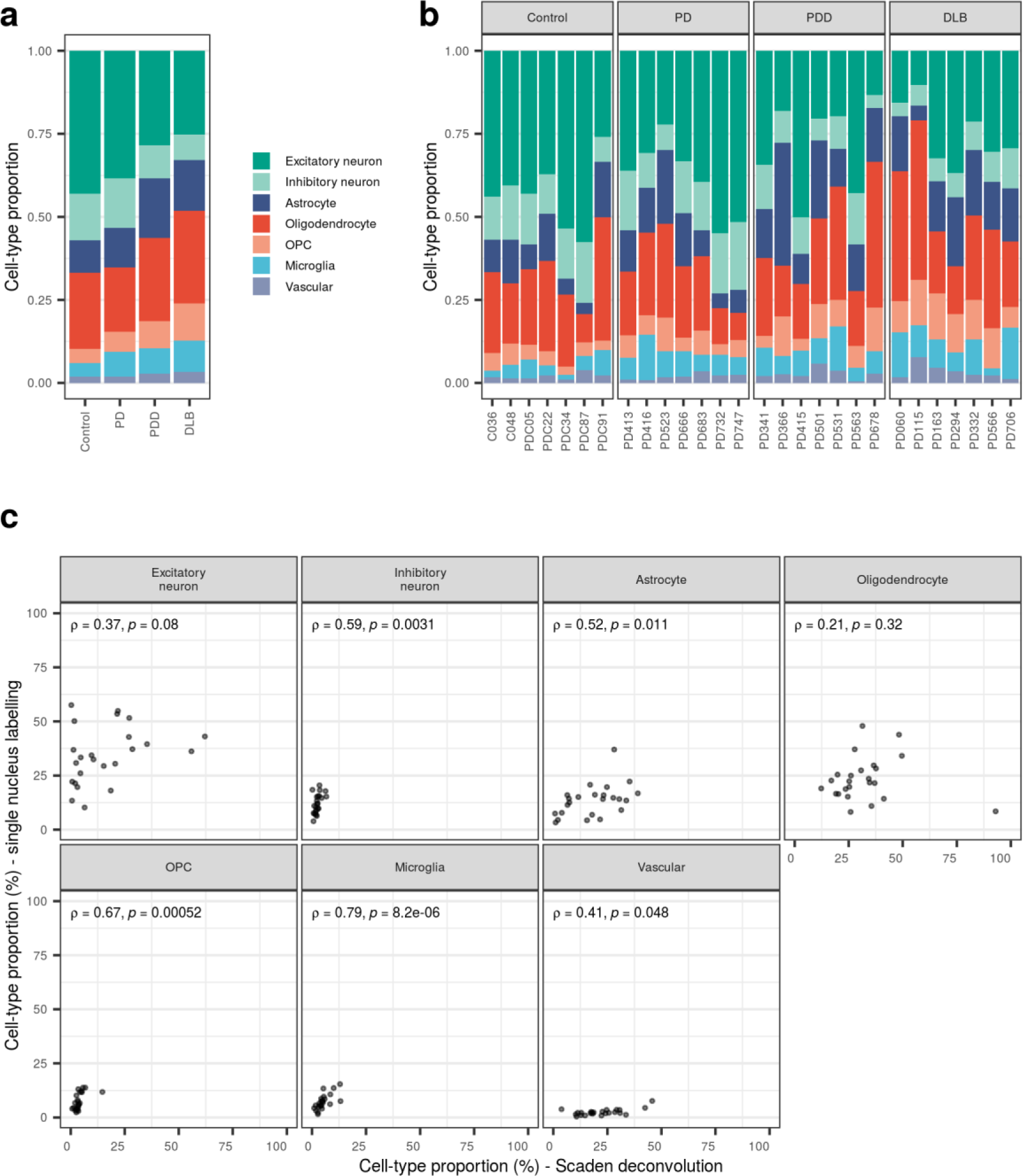
Consistency of cell types across individuals. **(a)** Proportion of nuclei of each cell type isolated across all individuals in each disease group (n = 7 per group) and **(b)** across each individual. **(c)** Scatterplot of cell-type proportions (%) derived from Scaden deconvolution and cell-type labelling of single nuclei. In each panel, Spearman’s rho (ρ) and associated p-value (p) are displayed. Proportions are available in **Supplementary Table 2**.

**Supplementary Figure 5.**
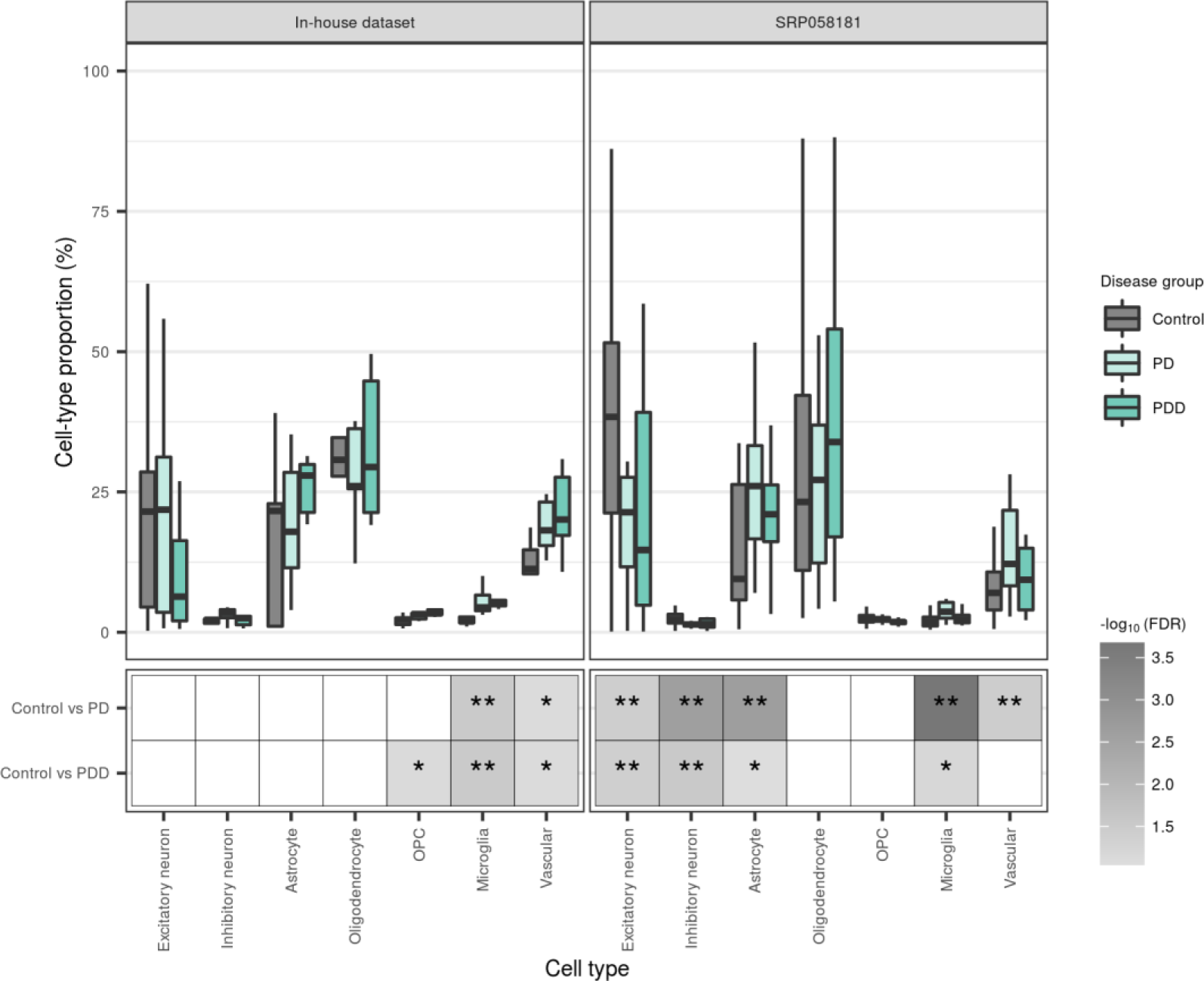
Replication of cell-type proportions. Cell-type proportions derived from Scaden deconvolution of our bulk-tissue RNA-sequencing or the recount dataset, SRP058181. Cell-type proportions (upper panel) are grouped by cell type and disease status and displayed relative to the median of controls (within a cell type). Significant differences in cell-type proportions between disease groups (lower panel) were determined using the Wilcoxon rank sum test, with FDR correction for multiple testing. Non-significant results (FDR > 0.1) were coloured white; **, FDR < 0.05; *, FDR <= 0.1. Results are available in **Supplementary Table 2**.

**Supplementary Figure 6.**
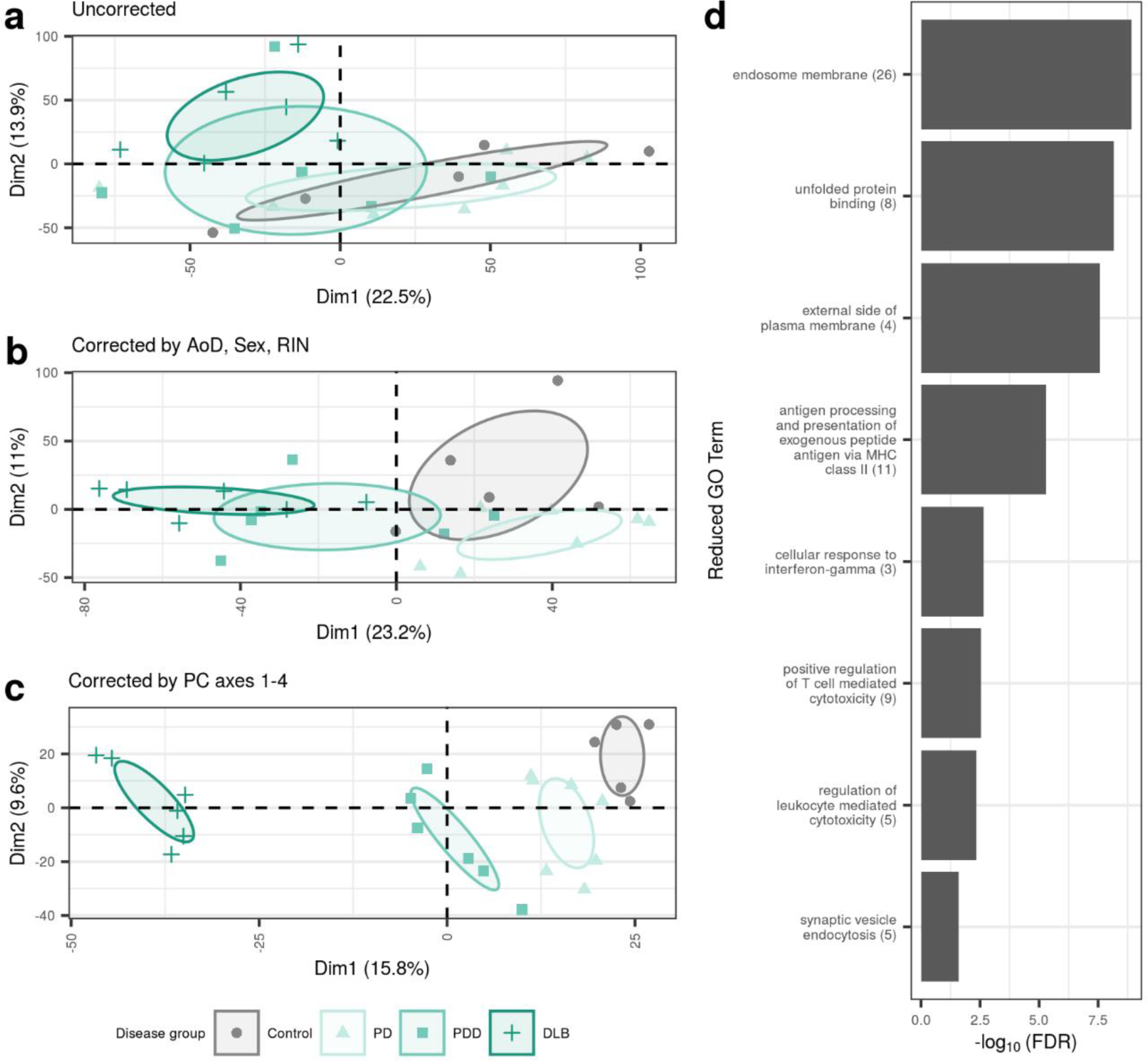
Effect of different batch-correction strategies on sample clustering by gene expression. Samples were plotted by the first two principal components derived from **(a)** uncorrected gene expression, **(b)** gene expression adjusted for age of death, sex and RIN and **(c)** gene expression adjusted for cell-type and experimental covariates. Principal component analyses were performed on gene-level expression filtered to include only genes with count > 0 in all samples (28,692 genes). Ellipses represent the 95% confidence level around group mean points (not displayed). **(d)** Reduced GO terms associated with the top 100 genes that contribute to PC1 in **(c)**. Original GO term enrichments (referred to as “child terms”) were grouped using semantic similarity. The number of enriched child GO terms assigned to each parent term is indicated in parentheses on the y-axis. Each parent term is represented by the most significant child term associated with it. Top 100 genes, their contributions to PC1 and results of pathway enrichment are available in **Supplementary Table 4**.

**Supplementary Figure 7.**
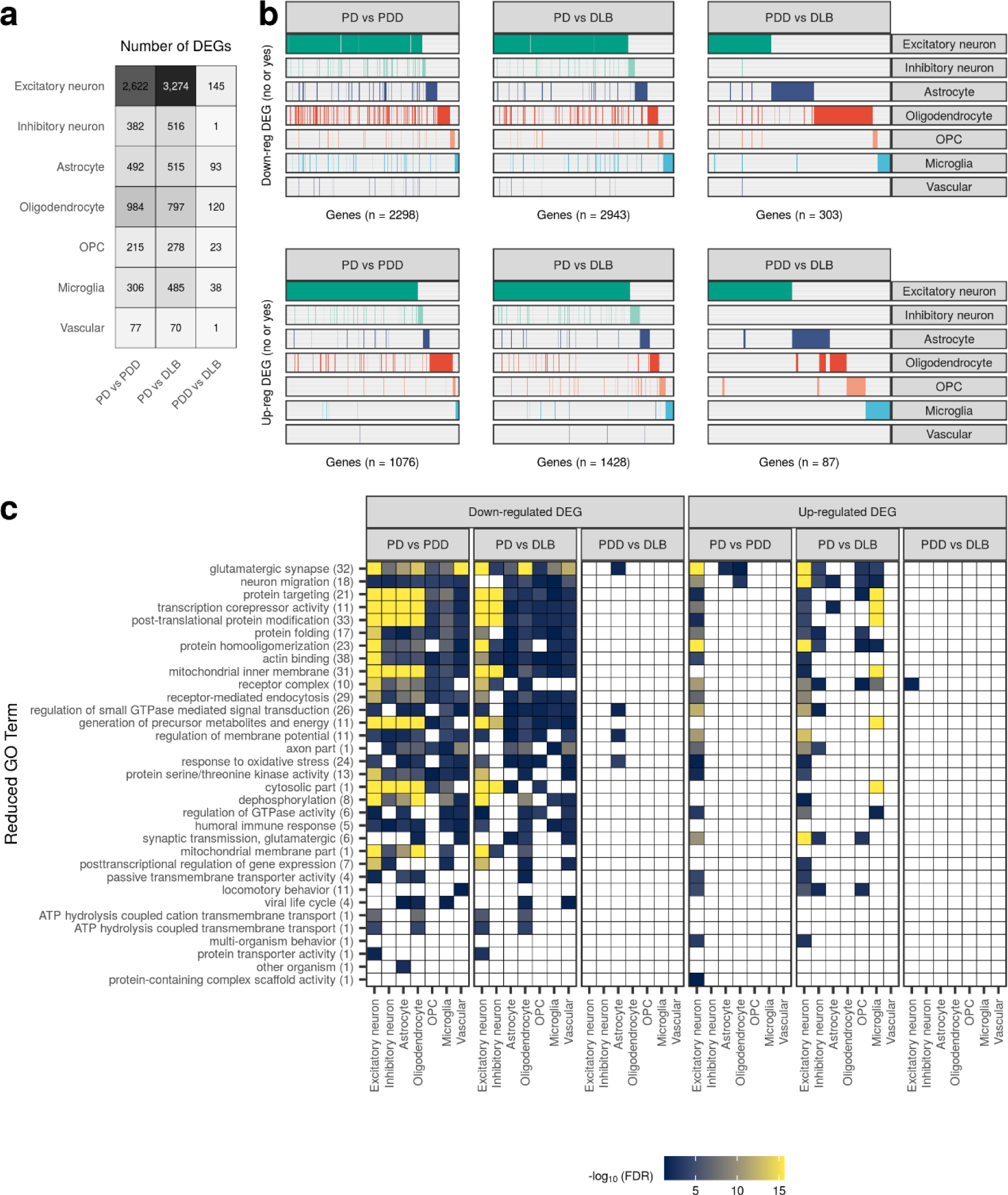
Cell-type-specific gene expression changes and pathway enrichments between disease states. **(a)** Number of differentially expressed (DE) genes (|log_2_(fold change)| ≥ log_2_(1.5), FDR < 0.05) across each cell type in pairwise comparisons between disease groups. The fill of the square is proportional to the number of DE genes. **(b)** Binary plot indicating with bars whether a gene (column) is down-regulated (upper panel) or up-regulated (lower panel) in a given cell type (rows). Number of DE genes in each comparison indicated on the x-axis. **(c)** Reduced GO terms associated with cell-type-specific down- and up-regulated DE genes identified across pairwise comparisons between disease groups. Due to the magnitude of pathway enrichments, original GO term enrichments (referred to as “child terms”) were grouped using semantic similarity. The number of enriched child GO terms assigned to each reduced parent term across all cell types and comparisons in the panel is indicated in parentheses on the y-axis. Reduced GO terms were ordered on the y-axis by the number of cell types and comparisons in which the term was found enriched. The fill of each tile indicates the - log_10_(FDR) of the most significant child term associated with the parent term within that comparison/cell type. Non-significant results (FDR > 0.05) were coloured white. All cell-type-specific DE genes and pathway enrichments are available in **Supplementary Table 5** and **Supplementary Table 6**, respectively.

**Supplementary Figure 8.**
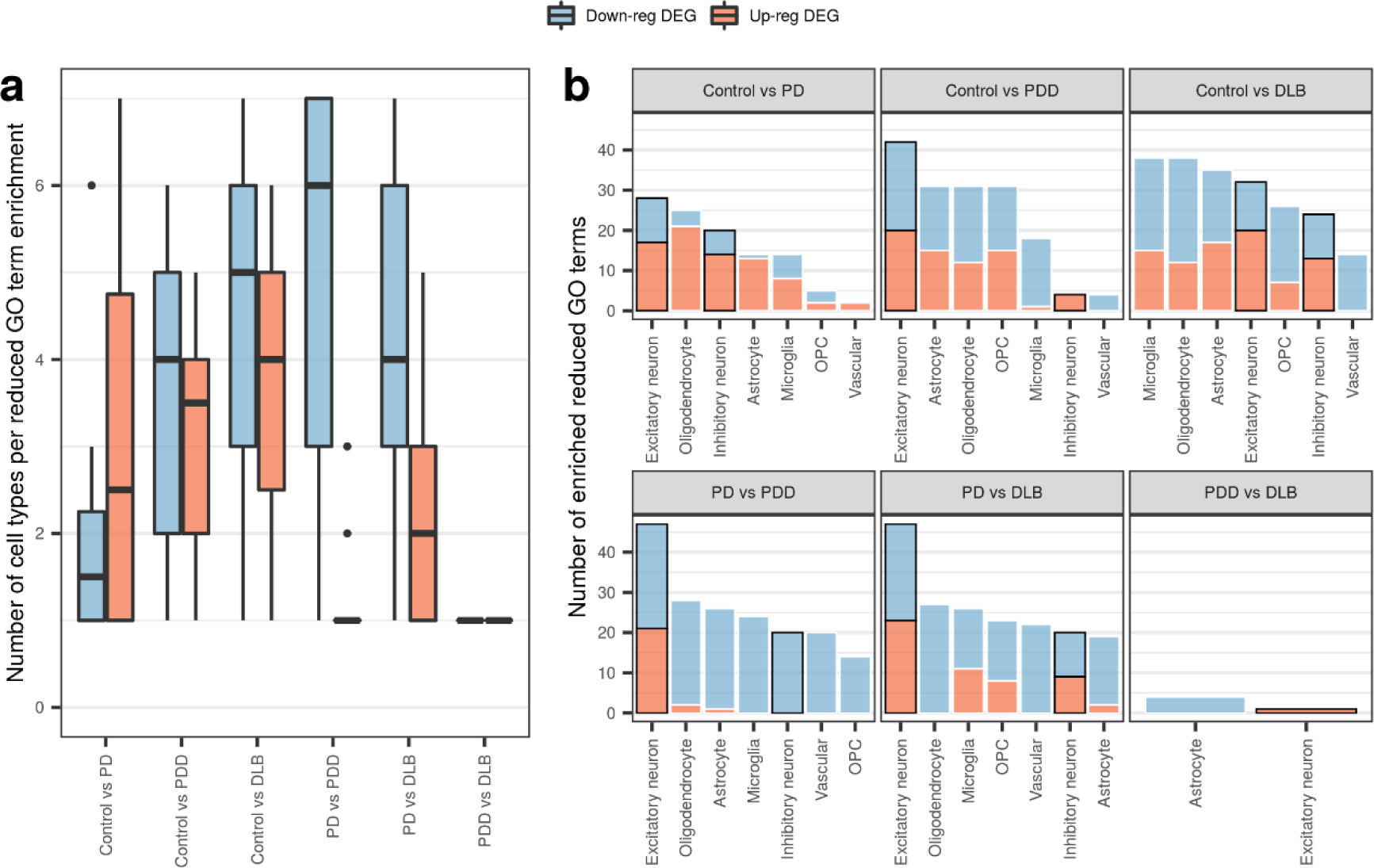
Reduced GO term counts across cell types and pairwise comparisons. **(a)** Number of cell types each GO term was associated with using cell-type-specific down- and up-regulated DE genes identified across pairwise comparisons. **(b)** Number of enriched GO terms identified using cell-type-specific down- and up-regulated DE genes from each pairwise comparison. Within each panel, cell types are ordered by the total number of enriched GO terms, irrespective of direction of effect, from highest to lowest. Excitatory and inhibitory neurons have been marked with a black border.

**Supplementary Figure 9.**
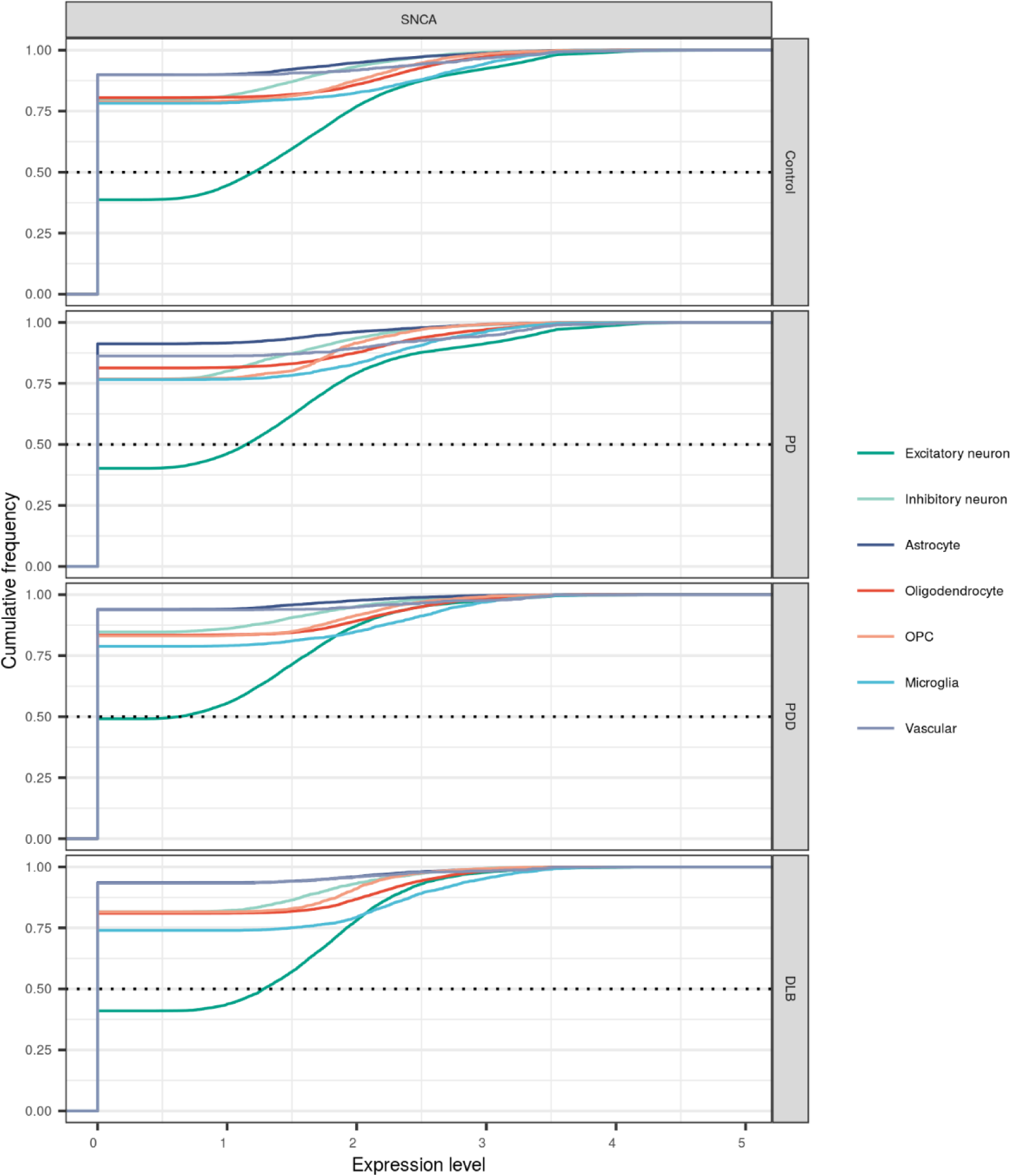
*SNCA* expression across cell types in each disease group. Cumulative distribution plot comparing *SNCA* expression levels in each disease group across cell types. Cumulative distribution plots display the proportion of data (y-axis) less than or equal to a specified value (x-axis). The horizontal dashed line denotes where 50% of the data lies.

**Supplementary Figure 10.**
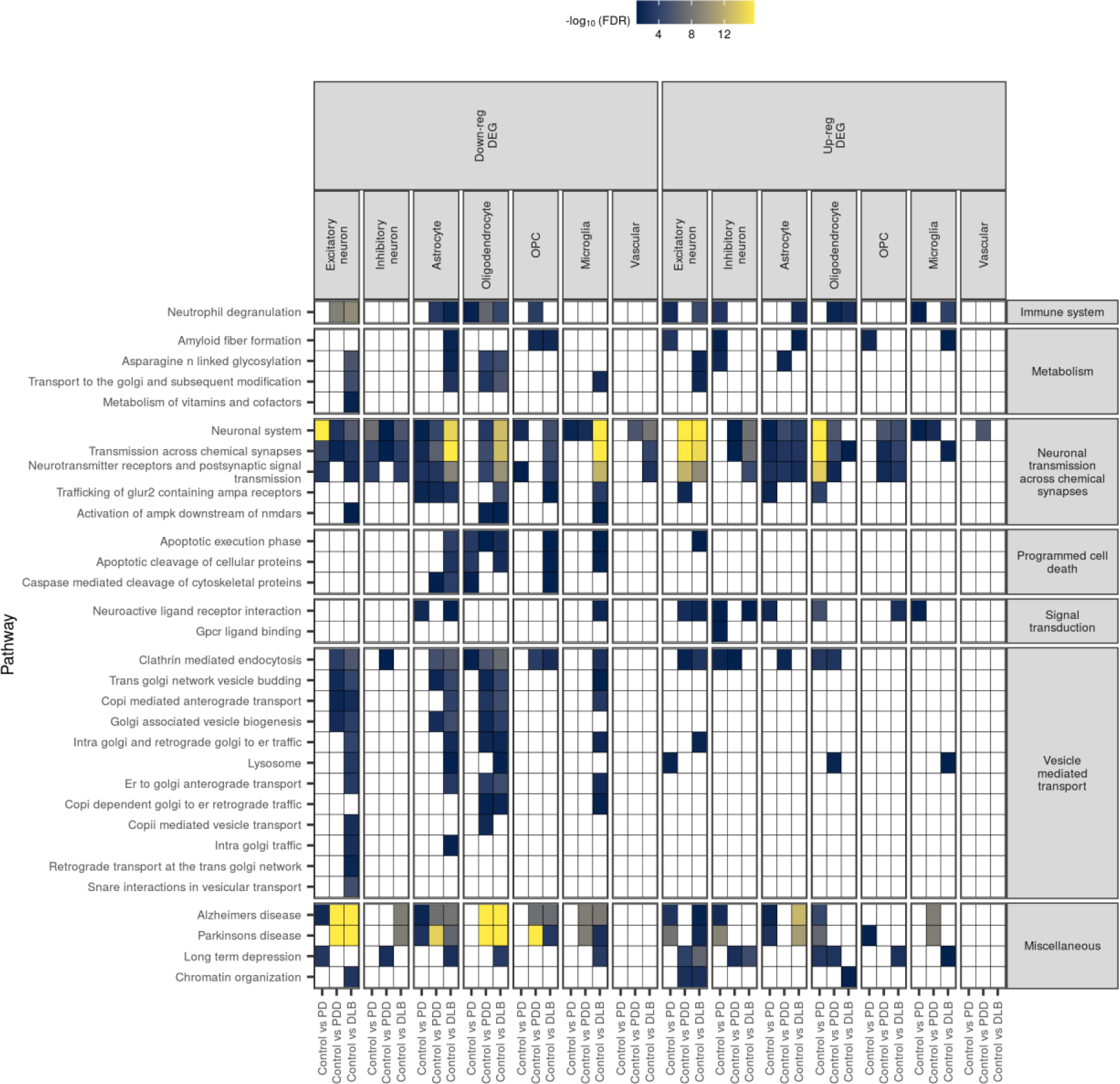
Cell-type-specific pathway enrichments across pathways genetically associated with PD. Pathway enrichments for all 46 PD-associated pathways (associated in a large-scale polygenic risk score-based assessment of 2,199 gene sets)[46]. The fill of each tile indicates the -log_10_(FDR) of enrichment. Non-significant results (FDR > 0.05) were coloured white. Pathway enrichment results are available in **Supplementary Table 7**.

**Supplementary Figure 11.**
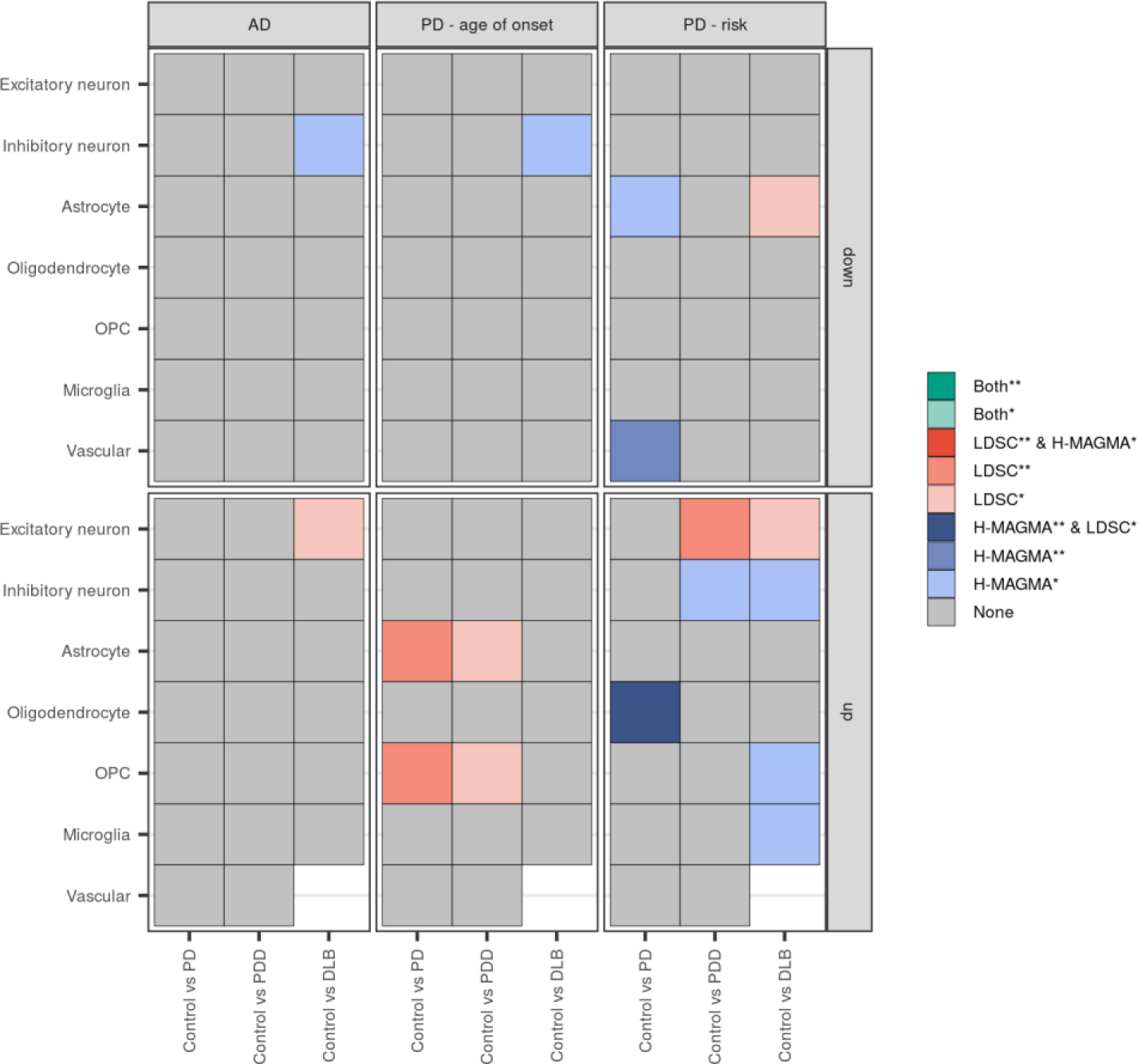
Genetic associations with cell-type-specific differentially expressed genes split by direction of effect. Genetic associations using down- and up-regulated cell-type-specific differentially expressed genes in disease comparisons with controls. Two methods were used to identify associations: Hi-C-coupled MAGMA (MAGMA) and stratified LD score regression (sLDSC). The heatmap is coloured by degree of significance with both or either method, with * and ** indicating nominal significance (unadjusted p-value < 0.05) or significance (FDR-corrected p-value < 0.05; corrected for number of cell types tested). Only 2 genes were up-regulated in vascular cells from Control vs DLB, thus no results were returned from either analysis. Results available in **Supplementary Table 8**. AD, Alzheimer’s disease; PD, Parkinson’s disease.

**Supplementary Figure 12.**
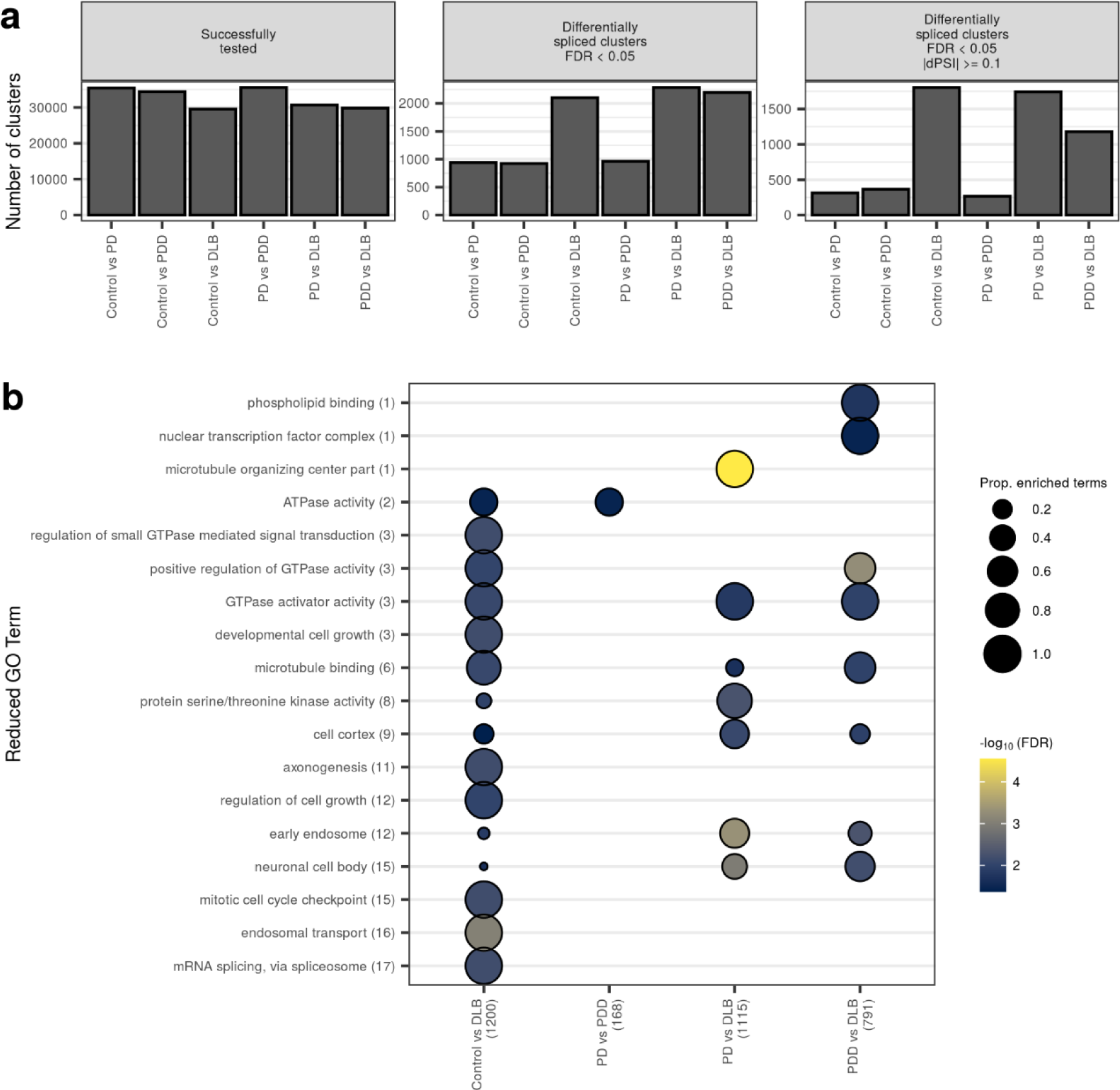
Differential splicing across disease groups. **(a)** Number of clusters that were successfully tested and found differentially spliced (DS) following correction for changes in cell-type proportions. **(b)** Reduced GO terms associated with genes found DS across pairwise comparisons (FDR < 0.05, |ΔPSI| >= 0.1). The number of DS genes is indicated in parentheses on the x-axis. Original GO term enrichments (referred to as “child terms”) were grouped using semantic similarity. The number of enriched child GO terms assigned to each parent term across pairwise comparisons is indicated in parentheses on the y-axis. Size of dot indicates the proportion of enriched child terms within a pairwise comparison, which is derived by dividing the number of enriched child terms by the total number of child terms assigned to a parent term. Fill of dot indicates the -log_10_(FDR) of the most significant child term associated with the parent term within that pairwise comparison. Results available in **Supplementary Table 11**.

**Supplementary Figure 13.**
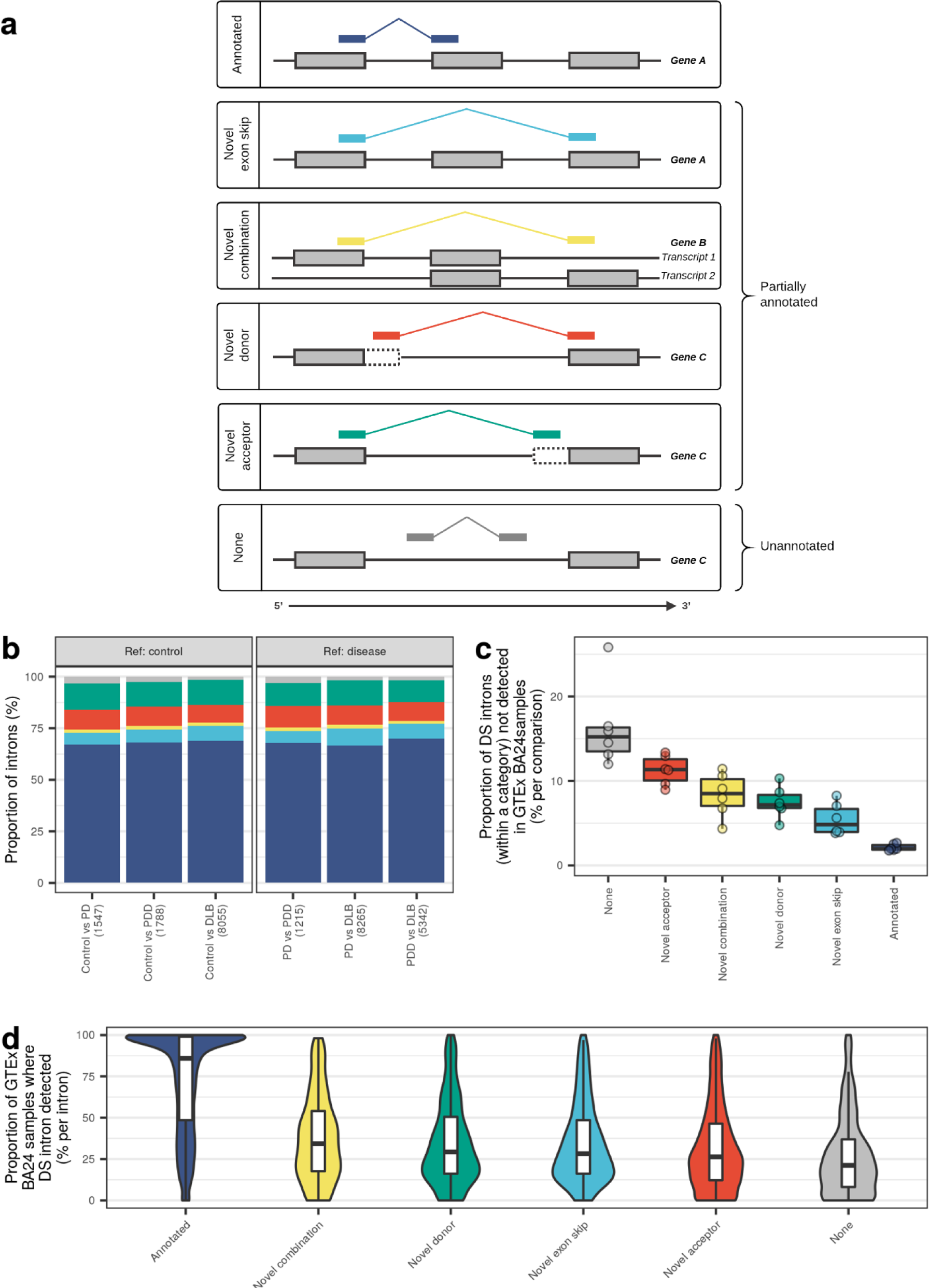
Annotation of differentially spliced introns. **(a)** Schematic illustration of the different categories of splicing event. Junction reads used to define Leafcutter introns were annotated based on their relationship to the annotated transcriptome (Ensembl v97). Here, the annotated transcriptome is illustrated by the grey-filled boxes. Annotated junctions have donor and acceptor splice sites that match the boundaries of an existing intron. Likewise, novel exon skip and novel combination junctions have donor and acceptor splice sites that overlap known exon boundaries derived from exons contained within the same transcript, but they represent introns which are not found in the set of annotated introns. They are distinguished by whether or not their donor and acceptor splice sites overlap exons derived from the same transcript. Novel donors and novel acceptors are junctions where only one end (3’ or 5’, respectively) matches the boundary of a known exon. All novel events are considered partially annotated. Unannotated junctions (“None”) have neither end overlapping a known exon. **(b)** Number of introns assigned to each category of splicing event as a proportion of all introns within the subset of differentially spliced intron clusters (FDR < 0.05, |ΔPSI| ≥ 0.1). The total number of junctions in each comparison is indicated in parentheses on the x-axis. **(c)** Number of DS introns that were undetected in a GTEx sample as a proportion of all DS introns within a category of splicing event. (**d**) Number of GTEx-derived anterior cingulate cortex (BA24) samples where a DS intron was detected as a proportion of the total number of BA24 samples (n = 99). Proportions in **(c)** and **(d)** are ordered by median from highest to lowest.

**Supplementary Figure 14.**
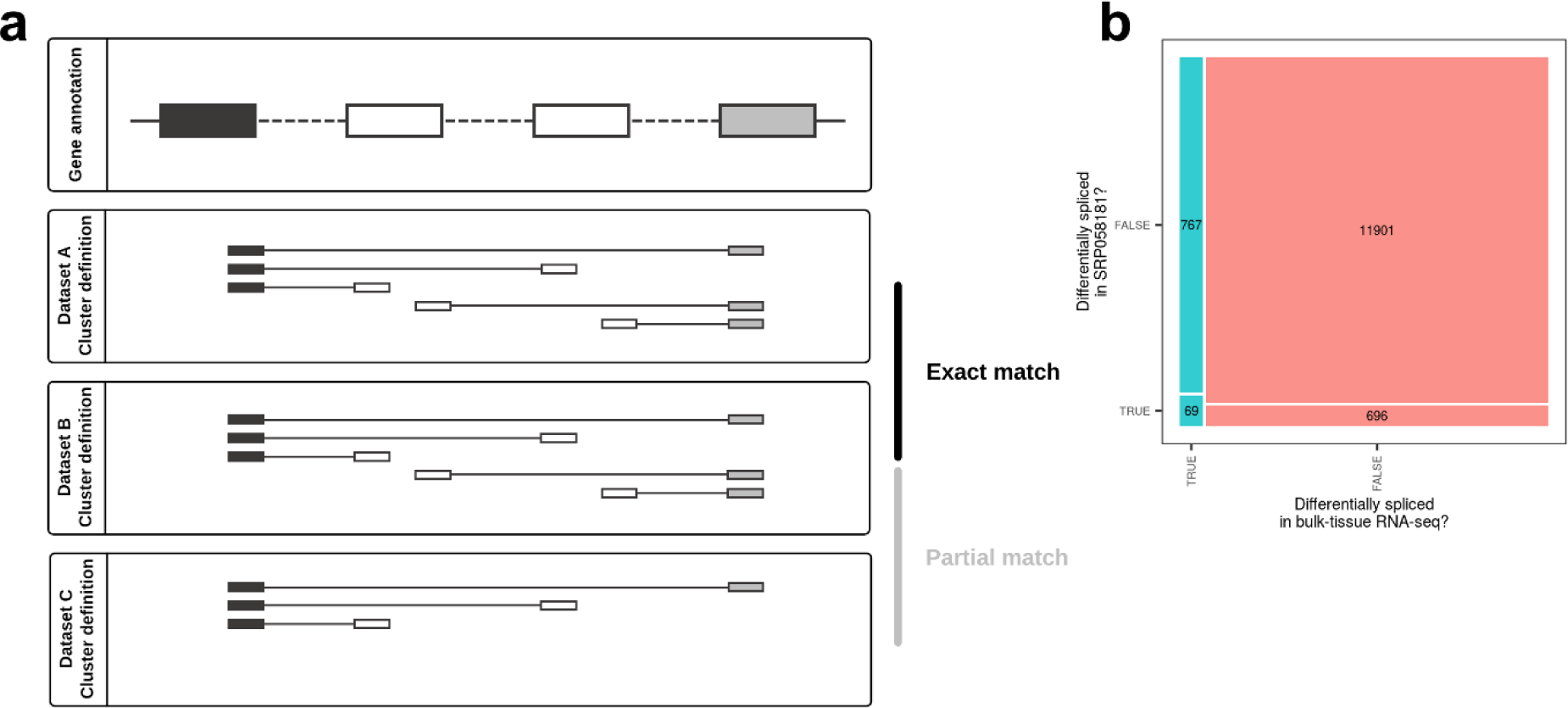
Defining replication across intron clusters and datasets. **(a)** Cluster definitions across datasets may vary; thus, comparisons of cluster definitions between datasets can yield exact, partial or no matches (no matches not illustrated). An exact match is defined as an intron cluster that contains the same introns, as determined by their splice donor and acceptor sites, across both datasets. In replication analyses, only exact matches between our dataset and the replication dataset from recount2 (recount ID: SRP058181) were carried forward. **(b)** Contingency table of differential splicing in our bulk-tissue RNA-sequencing and in the recount2 dataset, SRP058181. Only clusters that matched exactly across the two datasets were used to construct the contingency table. This yielded a total of 13,433 exactly matching intron clusters, 836 of which passed FDR < 0.05 in the discovery dataset. Unadjusted p-values in the replication dataset for these 836 overlapping clusters were then FDR-corrected, and any of the 836 that passed FDR < 0.05 in the replication dataset were considered validated. For gene-level analyses, only those validated intron clusters with ≥ 1 intron that shared the same direction of effect across both datasets were carried forward.

**Supplementary Figure 15.**
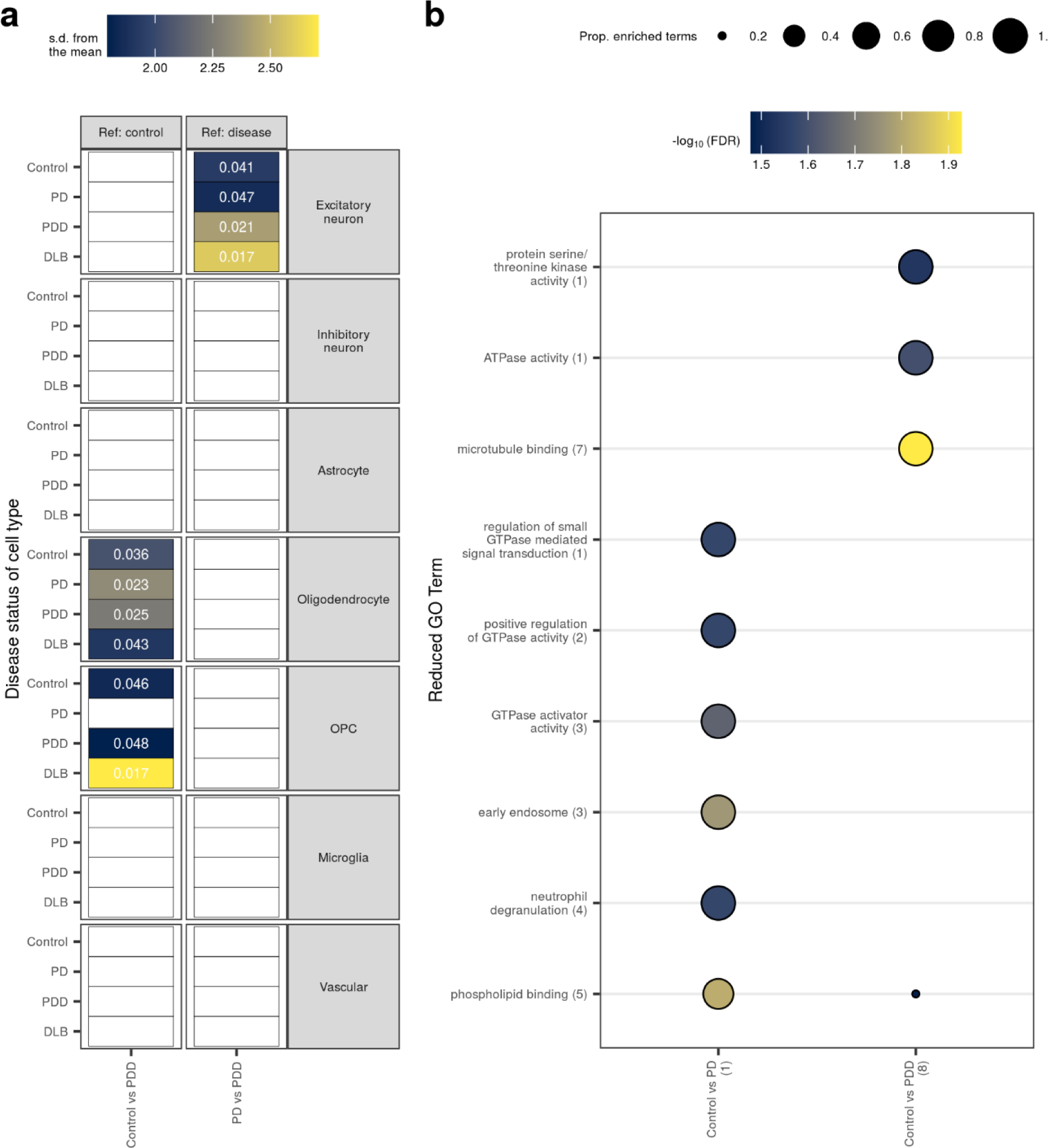
Cell-type and pathway enrichment analyses of validated differentially spliced genes. **(a)** Enrichment of validated differentially spliced (DS) genes in cell types derived from each disease group. Enrichments were determined using EWCE. DS at a gene-level was considered validated provided intron clusters matched exactly between our bulk-tissue RNA-sequencing and the SRP058181 dataset, passed FDR < 0.05 in both datasets, and at least one intron in the cluster shared the same direction of effect across both datasets. The x-axis denotes the groups compared in the differential splicing analysis, while the y-axis denotes the cell type and the disease status of specificity matrix from which it is derived. Standard deviations from the mean indicate the distance (in standard deviations) of the target list from the mean of the bootstrapped samples. No results survived FDR correction (FDR < 0.05); displayed are unadjusted p-values. Results with unadjusted p > 0.05 were coloured white. **(b)** Reduced GO terms associated with validated DS genes; the number of DS genes is indicated in parentheses on the x-axis. Original GO term enrichments (referred to as “child terms”) were grouped using semantic similarity. The number of enriched child GO terms assigned to each parent term across pairwise comparisons is indicated in parentheses on the y-axis. Size of dot indicates the proportion of enriched child terms within a pairwise comparison, which is derived by dividing the number of enriched child terms by the total number of child terms assigned to a parent term. Fill of dot indicates the -log_10_(FDR) of the most significant child term associated with the parent term within that pairwise comparison. Results of EWCE and pathway analyses are available in **Supplementary Table 10** and **Supplementary Table 11**, respectively.

**Supplementary Figure 16.**
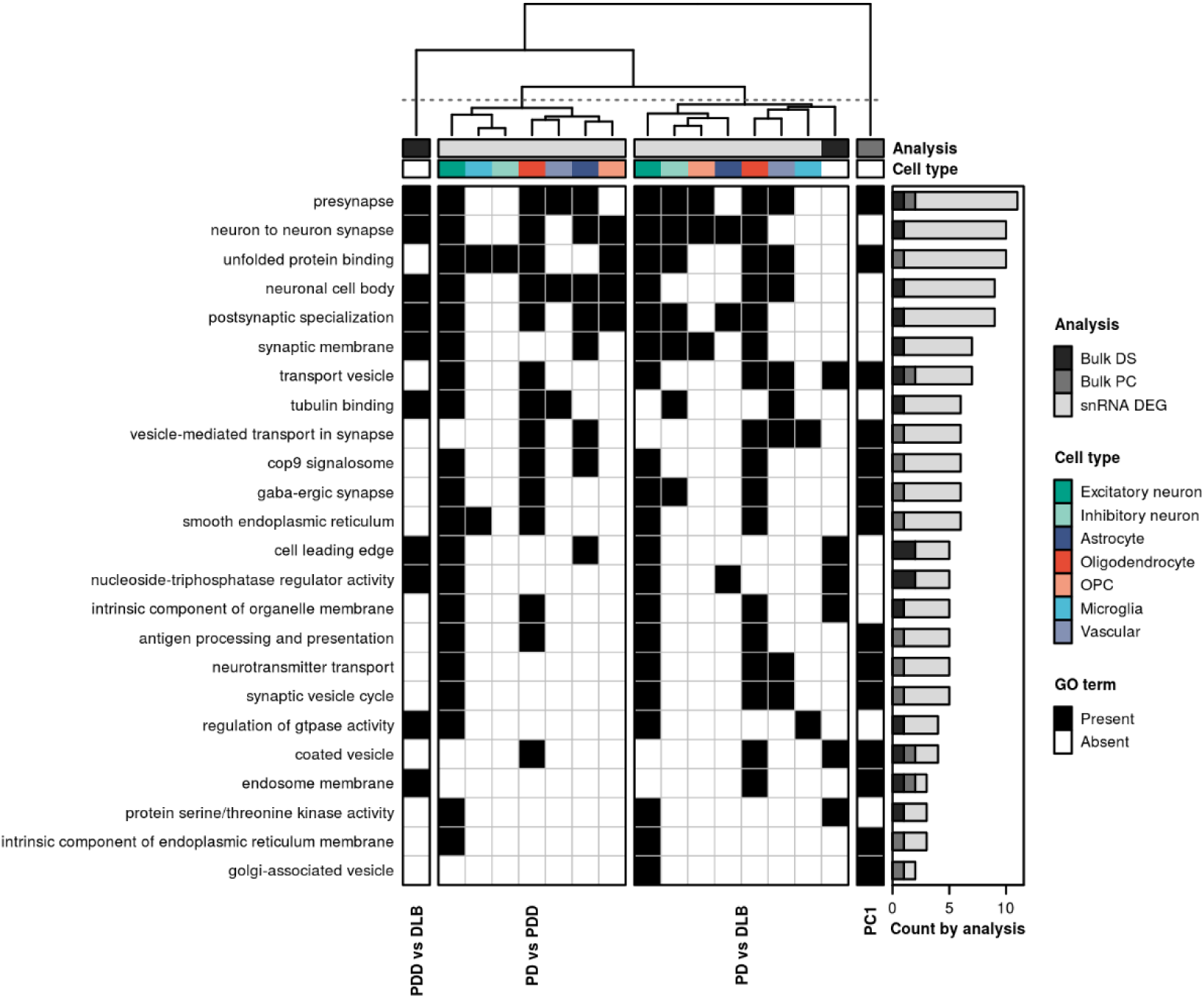
Pathway sharing between disease states. Clustering of shared pathway enrichments using genes identified across the three main analyses (represented by grey bar entitled, “Analysis”). These included: bulk-tissue differential splicing (“Bulk DS”, **Supplementary Figure 12**); gene contributions to bulk-tissue gene expression PC1 (“Bulk PC”, **Supplementary Figure 6**); and single-nucleus differential expression (“snRNA DEG”, **Figure 3**). Pathways (in rows) from all three analyses were filtered to include only those that appear across more than one type of analysis. Pathways are ordered from highest to lowest by the number of gene sets in which they are enriched (as displayed in the bar plot on the right-hand side). Gene sets (in columns) are clustered using hierarchical clustering on the Pearson correlation between gene sets (pathways were encoded with a binary 1 for “Present” or 0 for “Absent”, represented on the plot by black and white, respectively). Gene sets derived from differential splicing (Bulk DS) were collapsed across our own dataset and the validation dataset, resulting in one gene set (column) per pairwise comparison. Likewise, gene sets derived from up- and down-regulated single-nucleus DE gene sets were collapsed across cell types (represented by the coloured bar entitled, “Cell type”), such that each cell type was represented by a single column.

## Supplementary Tables

**Supplementary Table 1. Subject demographics, sample information, pathological measures and sequencing metrics.**

**Supplementary Table 2. Cell-type proportions from snRNA-sequencing and Scaden. Supplementary Table 3. List of bulk-tissue differentially expressed genes.**

**Supplementary Table 4. List of top 100 genes associated with gene expression PC1 following correction for cell-type proportions and experimental covariates.**

**Supplementary Table 5. List of cell-type-specific differentially expressed genes. Supplementary Table 6. List of cell-type-specific GO pathway enrichments.**

**Supplementary Table 7. List of cell-type-specific pathway enrichments using pathways genetically associated with PD.**

**Supplementary Table 8. H-MAGMA and sLDSC results. Supplementary Table 9. List of differentially spliced intron clusters.**

**Supplementary Table 10. EWCE results for differentially spliced genes. Supplementary Table 11. Pathway enrichments for differentially spliced genes.**

**Supplementary Table 12. List of RBPs binding motifs with a significant enrichment in differentially spliced proximal intronic regions compared with non-differentially spliced proximal intronic regions.**

## Supplementary Note

### Validation of differentially spliced introns in GTEx

#### Methods

Junction read counts from 99 GTEx-derived anterior cingulate cortex samples were accessed from recount2 (recount accession ID: SRP012682; GTEx v6) [95,108]. Paired-end 76-bp sequencing was applied to each sample, with a mean depth of 94.5 million read pairs per sample. All samples were of high quality with RIN values ranging from 5.5-8.9 and a median of 6.8. A DS intron was considered “detected in a GTEx sample” if its donor and acceptor splice sites precisely matched that of a junction with a count > 0 in a GTEx sample. Thereafter, two proportions were calculated: (i) the proportion of DS introns not detected in a single GTEx sample and (ii) the proportion of GTEx samples in which a DS intron was detected. The first proportion was calculated separately for each pairwise comparison by dividing the number of undetected DS introns by the total number of DS introns within a category of splicing event. The second proportion was calculated separately for each intron in a pairwise comparison by dividing the number of GTEx samples in which the intron was detected by the total number of GTEx samples.

#### Results

To determine whether DS introns were commonly observed in unaffected control tissue, a reference set of 99 control anterior cingulate cortex samples derived from the GTEx project was used. Across comparisons, between 3.8-5.4% of all DS introns went entirely undetected in GTEx samples. Detection rates varied across different categories of splicing event. Depending on the category of splicing event, anything between 1.8-26% of DS introns assigned to the category went entirely undetected in GTEx samples (**Supplementary Figure 13c**). This proportion was lowest for annotated and highest for unannotated categories, as might be expected under the assumption that an event that does not exist in the reference transcriptome remains unannotated by virtue of the low likelihood of detecting it. Of those DS introns that were detected in GTEx samples, 50% of annotated, partially annotated and unannotated events were observed in greater than 85.9%, 28.3%, and 21.2% of GTEx samples, respectively (**Supplementary Figure 13d**). Thus, despite the relatively high proportion of partially annotated DS introns, the ability to detect these events in larger control cohorts suggested these were biologically relevant splicing events.

#### Replication of differentially spliced introns in GTEx

##### Methods

Replication of differential splicing was performed using a replication dataset (see **Processing of PD case-control replication dataset**). Junction read counts were accessed from recount2, filtered to remove any regions that overlap ENCODE blacklist regions [92], and converted to .`junc` files. Intron clustering (which yielded 37,021 clusters encompassing 128,800 introns) and differential splicing were performed using the same parameters as above. As intron cluster definitions are dataset-dependent, only those intron clusters that matched exactly between the discovery and replication dataset were used for replication purposes. An exact match was defined as an intron cluster that contained the same introns, as determined by their splice donor and acceptor sites, across both datasets. This yielded a total of 13,433 exactly matching intron clusters, 836 of which passed FDR < 0.05 in the discovery dataset. Unadjusted p-values in the replication dataset for these 836 overlapping clusters were then FDR-corrected, and any of the 836 that passed FDR < 0.05 in the replication dataset were considered validated (**Supplementary Figure 14**). For gene-level analyses, only those validated intron clusters with ≥ 1 intron that shared the same direction of effect across both datasets were carried forward.

##### Results

Replication of differential splicing was performed using the same external PD case-control bulk-tissue RNA-sequencing dataset used in replication of deconvolution results. Only those intron clusters that were found to exactly match between datasets were used for replication (n = 13,433 intron clusters). Of these, 836 and 765 were DS (in at least one pairwise comparison) in our dataset and the replication dataset, respectively, with 69 shared between both (p-value = 0.001956; odds ratio = 1.53; 95% CI = 1.17-1.99; Fisher’s exact test; **Supplementary Figure 14**). We performed cell-type and pathway enrichments on genes containing a shared validated DS intron cluster with at least 1 intron with the same direction of effect in both datasets (*n* unique = 15 genes). Among cell-type enrichment tests, no gene sets passed FDR correction. However, nominally significant enrichments were observed in oligodendrocytes using genes found DS in PDD compared with control, similar to what we observed with our own dataset (**Supplementary Figure 15a**). Furthermore, several pathway enrichments observed were related to phospholipid binding, endosomes and GTPase activity (**Supplementary Figure 15b**).

